# How distributed subcortical integration of reward and threat may inform subsequent approach-avoidance decisions

**DOI:** 10.1101/2024.04.22.590530

**Authors:** Anneloes M. Hulsman, Felix H. Klaassen, Lycia D. de Voogd, Karin Roelofs, Floris Klumpers

**Author notes:** Correspondence (AMH), (FK). This work was funded by the Netherlands Organization for Scientific Research (NWO) Research Talent Grant #406-18-540 awarded to AMH and a European Research Council (ERC) Consolidator Grant #ERC_CoG-2017_772337 awarded to KR.

## Abstract

Healthy and successful living involves carefully navigating rewarding and threatening situations by balancing approach and avoidance behaviours. Excessive avoidance to evade potential threats often leads to forfeiting potential rewards. However, little is known about how reward and threat information is integrated neurally to inform approach or avoidance decisions. In this preregistered study, participants (N_behaviour_=31, 17F; N_MRI_=28, 15F) made approach-avoidance decisions under varying reward (monetary gains) and threat (electrical stimulations) prospects during functional MRI scanning. In contrast to theorized *parallel subcortical* processing of reward and threat information before cortical integration, Bayesian Multivariate Multilevel analyses revealed subcortical reward and threat *integration* prior to indicating approach-avoidance decisions. This integration occurred in the ventral striatum, thalamus, and bed nucleus of the stria terminalis (BNST). When reward was low, risk-diminishing avoidance decisions dominated, which was linked to more positive tracking of threat magnitude prior to indicating avoidance than approach decisions across these regions. In contrast, the amygdala exhibited dual sensitivity to reward and threat. While anticipating outcomes of risky approach decisions, we observed positive tracking of threat magnitude within the salience network (dorsal anterior cingulate cortex, thalamus, periaqueductal gray, BNST). Conversely, after risk-diminishing avoidance, characterized by reduced reward prospects, we observed more negative tracking of reward magnitude in the ventromedial prefrontal cortex and ventral striatum. These findings shed light on the temporal dynamics of approach-avoidance decision-making. Importantly, they demonstrate the role of subcortical integration of reward and threat information in balancing approach and avoidance, challenging theories positing predominantly separate subcortical processing prior to cortical integration.

## INTRODUCTION

Healthy and successful living involves carefully navigating rewarding and threatening situations by adopting an appropriate balance between approach and avoidance behaviours. Disruption of this balance can result in severe consequences, as observed in anxiety-related disorders and addiction (American Psychiatric Association, 2013). In approach-avoidance decision-making, the weighing of potential reward and threat prospects is pivotal (Corr, 2013). However, little is known about how this weighing is neurally represented to inform approach or avoidance decisions. In this study, we leveraged Bayesian Multivariate Multilevel (BMM) analyses on trial-by-trial BOLD fMRI data to investigate how distributed processing and integration of reward and threat information in the human brain contribute to approach-avoidance decision-making.

Human neuroimaging studies and recent theoretical frameworks suggest that reward and threat information is processed in distinct *parallel* subcortical streams, followed by reward-threat *integration* in cortical regions (Yacubian et al., 2006; Aupperle and Paulus, 2010; Basten et al., 2010; Park et al., 2011; Schlund et al., 2016; Zorowitz et al., 2019; Azab and Hayden, 2020; Livermore et al., 2021). For example, the ventral striatum (vStriatum) plays a crucial role in processing positive value and rewards (Talmi et al., 2009; Aupperle and Paulus, 2010; Basten et al., 2010; Bartra et al., 2013), potentially driving approach behaviour (Alvarez & Ruarte, 2001; Beaufour et al., 2001; Pedersen et al., 2021). Conversely, the amygdala and bed nucleus of the stria terminalis (BNST) are widely recognized for their pivotal roles in threat processing (Shackman and Fox, 2016; Abivardi et al., 2020; Hur et al., 2020; Hulsman et al., 2021c) and facilitation of avoidance behaviour (Ch’ng et al., 2018; Duque-Wilckens et al., 2018; Giardino et al., 2018; Miller et al., 2019; Hu et al., 2020). Value signals, indicating reward and threat potential, subsequently need to be integrated to inform decision-making, as previously demonstrated in cortical regions including the ventromedial prefrontal cortex (vmPFC) and dorsal anterior cingulate cortex (dACC) (Basten et al., 2010; Park et al., 2011; Schlund et al., 2016; Zorowitz et al., 2019; Azab and Hayden, 2020).

However, preclinical studies have demonstrated that reward and threat processing may be *intertwined* at the neuronal level in subcortical regions, with processing of opposite valence occurring in neighbouring cells, e.g., within amygdala subregions (Gentry et al., 2016; Burgos-Robles et al., 2017; Hamel et al., 2017; Beyeler et al., 2018). Moreover, recent rodent studies have shown that subcortical regions, including the central nucleus of the amygdala and nucleus accumbens can trigger both approach and avoidance motivation depending on the context and input received from other structures (Warlow et al., 2020; Zhou et al., 2022) and that regions of the thalamus respond particularly to situations requiring integration, such as approach-avoidance conflict (Engelke et al., 2021). This aligns with recent human work showing that regions of the salience network (thalamus, periaqueductal gray, dACC, and anterior insula) are sensitive to both rewards and threats (Aupperle et al., 2015; Schlund et al., 2016; Moughrabi et al., 2022). Strikingly, however, none of the aforementioned human studies investigated how the interplay between reward and threat is represented across these brain regions and how this balance shifts upon decisions to approach or avoid.

To investigate this, participants performed a Fearful Avoidance Task (Hulsman et al., 2021a) in the MRI scanner. In this task, participants made approach-avoidance decisions under varying prospects of reward (monetary gain) and threat (electrical shock). Based on previous human literature, we hypothesized that prior to approach, activations would occur in brain regions most strongly associated with appetitive processing (vStriatum, vmPFC). Conversely, prior to avoidance, we expected activations in brain regions traditionally linked to defensive responding (amygdala, BNST). Additionally, we expected that the vmPFC and dACC would play a crucial role in reward-threat integration. Finally, following risky approach decisions, leading to an increased chance of receiving shocks or rewards, we expected increased activity in regions associated with appetitive (vStriatum, vmPFC), defensive (amygdala, BNST), and salience processing (thalamus, periaqueductal gray, dACC, anterior insula) compared to risk-diminishing avoidance.

## MATERIALS AND METHODS

Research questions, hypotheses, and planned analyses were preregistered at the Open Science Framework: https://osf.io/7sm9k (Hulsman et al., 2021b) prior to data analyses. All data and code will be available at the Donders Institute for Brain Cognition and Behaviour repository upon publication: https://doi.org/10.34973/gxr8-9f24. Supplementary information is available at the Open Science Framework: https://osf.io/hbjp5.

### Participants

The recruited sample consisted of 31 participants (17 females, *M*_age_ = 23.45, *SD*_age_ = 3.33). Participants had an average trait anxiety score of *M*_STAI-T_ = 38.03 (*SD* = 8.39, range = 27-55; one participant did not complete the State-Trait Anxiety Inventory (STAI; Spielberger et al., 1970). Three participants were a priori excluded from the MRI analyses due to a lack of avoidance behaviour (<10%, see also Hulsman et al., 2024). As a result, planned contrasts of approach vs. avoid trials could not be carried out in these participants. Exclusion of these participants led to a final sample size of N=28 (15 females, *M*_age_ = 23.25, *SD*_age_ = 3.27). Participants self-reported meeting the following inclusion criteria: between 18 and 35 years of age, normal/corrected-to-normal vision, sufficient comprehension of English or Dutch, no psychiatric disorders, no neurological or cardiological disease/treatment, no use of psychotropic medication, no current drug or alcohol abuse, and no epilepsy. Participants received EUR 10 for participation and an additional bonus between EUR 0-5 depending on their performance in the Fearful Avoidance Task (described below). All participants were MRI eligible and provided written informed consent. This study was carried out in compliance with the declaration of Helsinki and approved by the local ethics committee (CMO Arnhem-Nijmegen).

### Procedure

Participants were informed that they had the opportunity to earn a monetary bonus based on their behaviour in the Fearful Avoidance Task (FAT, Hulsman et al., 2021a). They were asked to write down 10 random numbers, which would later be linked to specific trial numbers using a mathematical formula that was disclosed to the participants only after the experiment. If they received a monetary reward during these trials, they would receive the corresponding sum as a bonus, capped at a maximum of EUR 5.

Next, participants underwent titration procedures to determine their individual reward and threat levels for the Fearful Avoidance Task. Initially, to establish shock intensity at a point of maximum discomfort without causing pain, participants underwent a standardized *shock work-up procedure* consisting of five shock administrations (Klumpers et al., 2010). Each shock was rated on a scale ranging from 1 (not painful/annoying) to 5 (very painful/annoying). Intensities were adjusted gradually to attain a subjective level that as maximally uncomfortable without being painful (aiming for a 4 on the 5-point scale). Electric shocks were delivered via two Ag/AgCl electrodes attached to the distal phalanges of the middle and ring finger of the right hand using a MAXTENS 2000 (Bio-Protech) device. Shock duration was 250ms at 150Hz, and intensity varied in 10 intensity steps between 0-80mA. We opted to use electric shocks in this paradigm because primary reinforcers may evoke stronger psychophysiological responses than for example affective pictures (Lissek et al., 2007; Delgado et al., 2011).

Next, we conducted a *reward-threat titration procedure* to determine the monetary reward required for individuals to accept the risk of receiving an electric shock of the predetermined shock work-up intensity. Amounts between EUR 0.20 and EUR 10 were presented in semi-random order. For each amount, participants were asked whether they would be willing to risk receiving electrical stimulation. No actual shock reinforcement was administered throughout this entire procedure. Participant’s indifference point (*M* = 1.17, *SD* = 1.84) was subsequently used to calculate the reward values for the ensuing FAT (described below).

Finally, the Fearful Avoidance Task (FAT, Hulsman et al., 2021a) was employed to evaluate active approach-avoidance behaviour under competing reward (monetary gains) and threat (electric shocks) prospects. In this study, participants performed an adapted version of the FAT (see next section), optimized for use in the MRI scanner.

### Fearful Avoidance Task (FAT)

Participants received on-screen instructions for the task in the MRI scanner. Each trial consisted of a cascade of three phases (see Figure 1A). In the *decision phase* participants were confronted with a combination of reward and threat. The reward level was a monetary amount displayed in numbers below the avatar and was relative to the individual’s indifference point (IP) either low (IP-40-50%), medium (IP ± 5%), or high (IP + 40-50%). The threat levels were displayed with different avatars (Figure 1B). In total, there were three threat levels associated with different durations of electrical shock (low: 5ms, medium: 35ms, and high: 250ms). Reward and threat were combined in a 3 x 3 full factorial manner, resulting in nine combinations of reward and threat. After trial onset, participants had 2.5 seconds to decide to approach or avoid by pulling the joystick towards themselves or pushing the joystick away from themselves, respectively. As confirmation of a timely response (<2.5s) a white line appeared below the reward for the remainder of the decision phase and the subsequent outcome anticipation phase.

**Figure 1.**
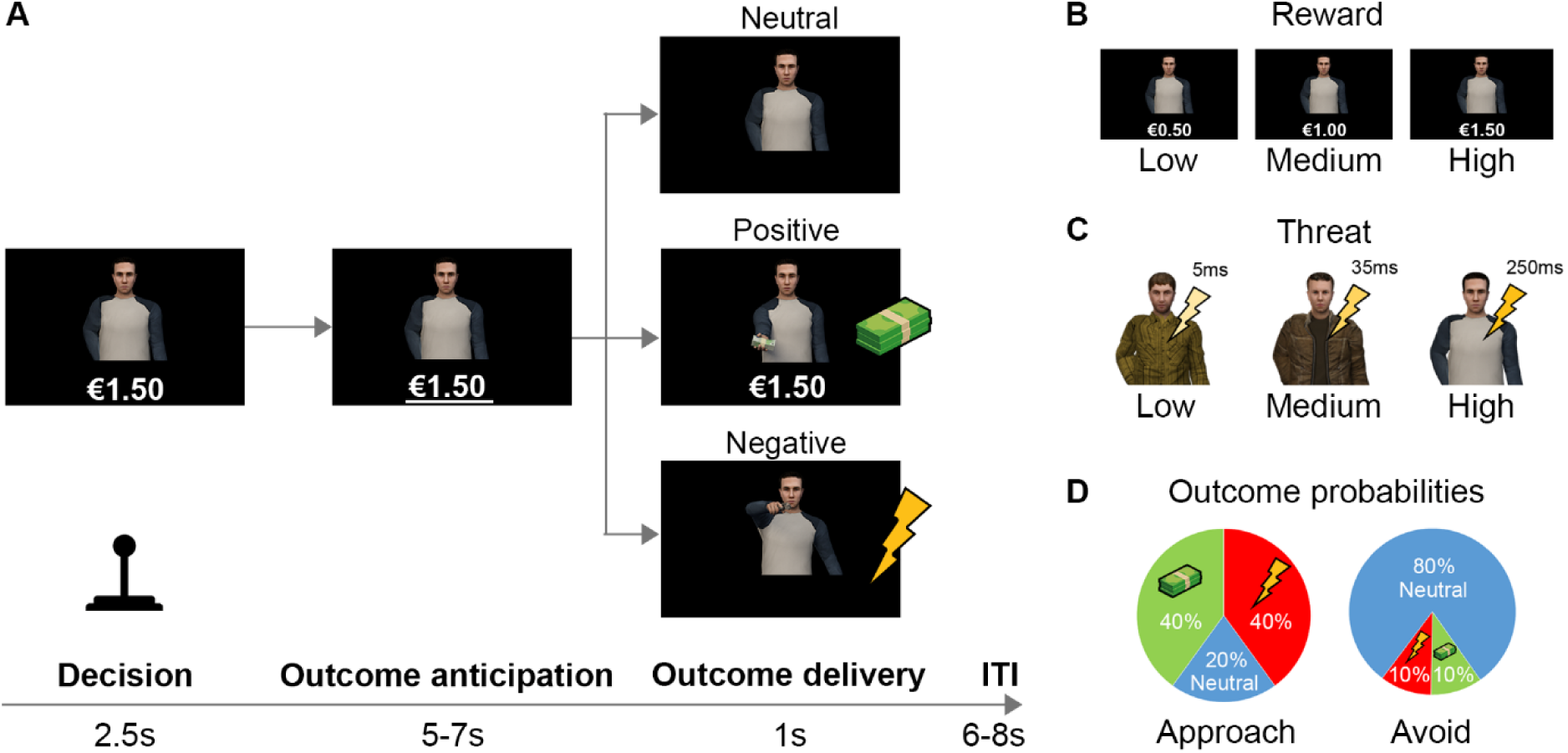
Overview of the Fearful Avoidance Task (FAT). **(A)** Example of the FAT trial structure. In each trial, participants were first confronted with a combination of reward (displayed in text) and threat (signalled by the avatar). Participants had 2.5s to decide to approach or avoid by pulling the joystick towards themselves or pushing it away, respectively. Next, they anticipated the outcome of their decision. Finally, the outcome of their decision was displayed. After receiving the outcome, a fixation cross was presented during the inter-trial interval (ITI). **(B)** Overview of the reward levels: reward levels were low (IP – 40-50%), medium (IP ± 5%), or high (IP + 40-50%), relative to the individual’s indifference point (IP). In this example, with an IP of €1.00, the low reward level ranged from €0.50 to €0.60, the medium reward level ranged from €0.95 to €1.05, and the high reward level ranged from €1.40 to €1.50. **(C)** Overview of the threat levels: three distinct avatars signalled shock duration. Contingencies between avatars and threat levels were counterbalanced across participants. **(D)** Overview of the outcome probabilities for approach and avoidance decisions. Approach decisions involved a high risk of receiving electric shocks or earning monetary rewards, while avoidance decisions diminished the risk of electric shocks but also reduced the potential for monetary rewards.

Next, a 5-7s *outcome anticipation phase* started. In this phase, the avatar and monetary amount remained on the screen while participants anticipated the outcome of their decision.

Finally, participants received one of three possible outcomes: positive, negative, or neutral. In a positive outcome, the avatar offered a stack of banknotes. In a negative outcome, the avatar drew a gun and shot. Additionally, participants received an electric shock with a duration that corresponded to the threat level symbolized by the avatar. During a neutral outcome, the avatar kept its hands behind its back, resulting in omission of both reward and threat. The probability of receiving a particular outcome depended on the decision that participants made (Figure 1C). Approach led to a 40/40% chance of receiving a positive or negative outcome and a 20% chance on receiving a neutral outcome, whereas avoidance led to an 80% chance of receiving a neutral outcome and a 10/10% chance of receiving a positive or negative outcome. These differential outcome contingencies of approach and avoidance were modelled after real-life approach-avoidance conflicts. In such conflicts, approach decisions typically entail a higher risk of encountering perceived threats and rewards, while avoidance decisions mitigate these threats but also forfeit potential rewards (Krypotos et al., 2018). Participants were instructed that late responses (>2.5s) always led to the negative outcome, i.e., getting shot and receiving an electrical shock of a duration that was symbolized by the avatar. After participants received the outcome, a fixation cross was displayed during the inter-trial interval (6-8s).

Since this study did not aim to investigate de novo contingency learning, participants received explicit instructions about the association between the avatar and its corresponding threat level, as well as the outcome probabilities of approach and avoidance decisions. Providing explicit instructions has been shown to significantly enhance learning of contingencies (Mertens et al., 2018), leading to stronger task effects, including those observed in neuroimaging measures (Mechias et al., 2010). Thus, contingencies were all clearly and repeatedly explained to the participants before starting the task, and comprehension was verified during a short practice session of 9 trials in the MRI scanner. The use of avatars as conditioned stimuli was intended to make the task more engaging. Previous research has shown that such stimuli can enhance motivation (Sailer et al., 2017) and sustain attention (Krath et al., 2021). The association between avatars and threat levels was counterbalanced across participants. In total, participants completed 90 trials (10 trials for each combination of reward and threat).

### MRI acquisition

Data was acquired on a 3T MAGNETOM PrismaFit MR scanner (Siemens AG, Healthcare Sector, Erlangen, Germany) using a 32-channel head coil. A T1-weighted scan was acquired in the sagittal orientation using a 3D MPRAGE sequence (TR=2000ms, TE=3.03ms, 192 sagittal slices; 1.0mm isotropic voxels; FOV=256mm). T2-weighted volumes were acquired using a multi-band multi-echo (MB3ME3) sequence, a fast sequence designed for whole brain coverage with reduced artefacts and signal dropout in medial prefrontal and subcortical regions (Cohen et al., 2018; Fazal et al., 2023) (TR = 1500ms, TE_1-3_ = 13.4/34.8/56.2ms, flip angle = 75°, 51 sagittal slices; 2.5mm isotropic voxels).

### Behavioural analyses

First, we verified whether the data adhered to the preregistered inclusion criteria, which stipulated a maximum of 50% missing responses and absence of atypical response patterns as defined by Hulsman, Klaassen, et al., (2021). Atypical response patterns refer to behaviour indicating that the participant may not have comprehended the task correctly or did not perform the task seriously (e.g., when participants avoided substantially more on low threat trials than high threat trials, see supplement for a detailed description of atypical response patterns). All participants met these inclusion criteria.

To investigate the effect of reward, threat, and their interaction on the decision (approach/avoid) that participants made, we ran a Bayesian multilevel model. For full model specifications, see equation (1) below. Other model specifications were as described in the next section.

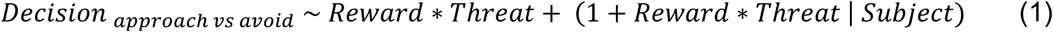

### Bayesian multilevel analyses (general)

As preregistered (Hulsman et al., 2021b), throughout behavioural and fMRI analyses, we employed linear Bayesian multilevel models. These linear Bayesian multilevel models were executed using R (Version 4.2.1; R Core team, 2022) within RStudio (Version 2022.12.0; RStudio Inc., 2009-2022) using the brms package (2.18.0, Bürkner, 2013 and Carpenter et al., 2017). All Bayesian multilevel models adhered to the following configurations: continuous predictors were standardized, and categorical predictors were coded using sum-to-zero contrasts. All models included a maximal random effects structure, consisting of a random intercept for each participant along with random slopes for all predictors and their interactions. Models with a binomial dependent variable (i.e., decision: approach/avoid) were modelled using a Bernoulli distribution, whereas models with continuous dependent variables (i.e., fMRI beta values) were modelled using a Gaussian distribution. Models were fitted using 4 chains with 4000 iterations each (2000 warm-up). A coefficient was deemed statistically significant when the associated ≥95% posterior credible interval was non-overlapping with zero. As recommended for analyses with an effective sample size <10.000 (Kruschke, 2014) this was supplemented with 90% posterior credible intervals when the 95% intervals were overlapping with 0, given that these intervals may yield more stable results (Makowski et al., 2019). These latter results are reported as trends when the 90% interval was non-overlapping with zero. When effects were non-overlapping with zero at the 90 or 95% interval, we additionally tested 99% and 99.9% intervals to further explore the likelihood of these effects (akin to reporting *p*<.01 and *p*<.001, respectively, rather than only *p*<.05).

### fMRI analyses

The fMRI data were processed using SPM12 (Wellcome Trust Centre for Neuroimaging, London, UK) following the standard preprocessing procedures. Functional scans were combined into a single image using the PAID-weighting method (Poser et al., 2006), co-registered to the anatomical scan, and normalized to the Montreal Neurological Institute (MNI) 152 T1-template using nonlinear transformations based on SPM’s tissue probability map. The normalized images (2mm isotropic) were then smoothed with an isotropic 3D Gaussian kernel with 6mm full width at half maximum (FWHM). For all models, high-pass filtering at 1 /128 Hz and a first-order autoregressive model were used as standard.

Our preregistered regions of interest (ROIs, see Figure 2) included the anterior insula (Shirer et al., 2012), amygdala (Rolls et al., 2020), bed nucleus of the stria terminalis (Avery et al., 2014), dorsal anterior cingulate cortex (Shirer et al., 2012), periaqueductal gray (Lojowska et al., 2015), thalamus (Rolls et al., 2020), ventral medial prefrontal cortex (Rolls et al., 2020), and ventral striatum (Piray et al., 2017).

**Figure 2.**
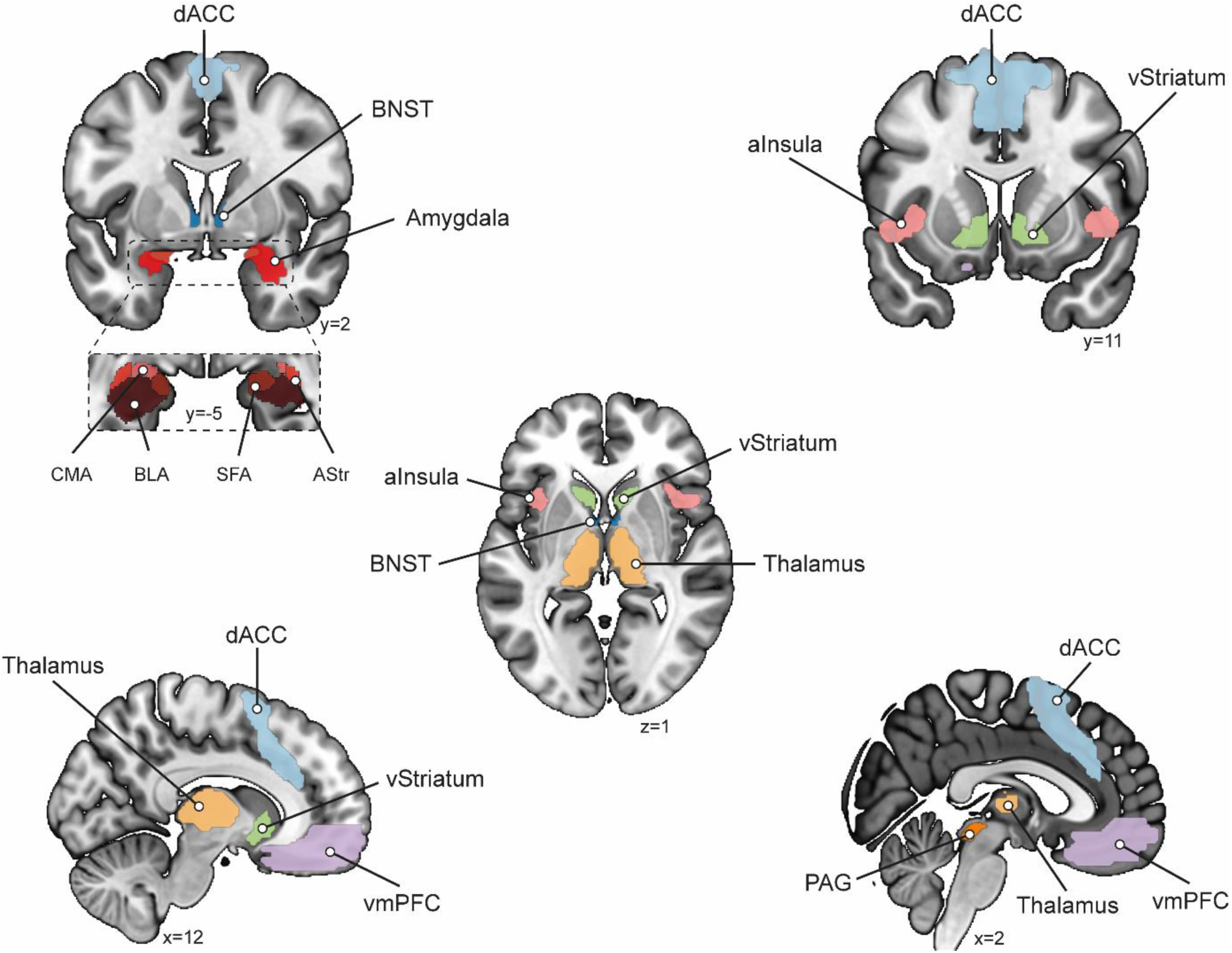
Regions of interest (ROIs). AInsula = anterior insula, amygdala, BNST = bed nucleus of the stria terminalis, dACC = dorsal anterior cingulate cortex, PAG = periaqueductal gray, thalamus, vmPFC = ventral medial prefrontal cortex, vStriatum = ventral striatum, CMA = central medial amygdala, BLA = basolateral amygdala, SFA = superficial amygdala, AStr = amygdalostriatal region. We hypothesized that the aInsula, dACC, PAG, and thalamus show sensitivity to both reward and threat. Additionally, we hypothesized that the amygdala and BNST are predominantly sensitive to threat, while the vmPFC and vStriatum are predominantly sensitive to reward. Furthermore, we expected that the dACC and vmPFC are involved in the integration of reward and threat.

To gain global insight into the neural activation patterns probed by the task, independent of the level of reward, the level of threat, and the decision, we first report the results of the whole-brain voxel-wise analysis contrasting neural activity during the *decision phase* against the baseline (i.e., the intertrial interval). All fMRI figures were created in MRIcroGL, with the use of the MNI 152 template image. As preregistered, whole-brain results were familywise (FWE) corrected for multiple comparisons according to random field theory at the cluster level (*p*<.05) after an initial cluster-defining threshold of *p*<.001 uncorrected.

### Bayesian Multivariate Multilevel (BMM) fMRI analysis: effects of reward, threat, and decision

To investigate the effects of reward, threat, decision, and their interactions on neural activity within our regions of interest, we leveraged Bayesian Multivariate Multilevel (BMM) analyses. In our task, we expected that approach and avoidance decisions would depend on specific task conditions, and indeed, in some individuals, conditions and behaviour were strongly correlated. For instance, in the low reward/high threat condition, several participants consistently decided to avoid. Given these circumstances, conventional MRI analyses are less suited to accurately disentangle the effect of reward/threat from the effect of the decision to approach or avoid). Particularly for these intricate designs, BMM analyses surpass conventional MRI analyses by effectively managing data dependency and unbalanced data, leading to increased accuracy of results (Chen et al., 2019a, 2019b). Therefore, BMM analyses emerged as the most optimal choice to investigate the effects of reward, threat, decision, and their interactions. Given that conducting BMM analyses on whole-brain voxel-wise level is not computationally feasible, we performed the BMM analyses on our preregistered regions of interest. In line with our preregistration, we also conducted two additional whole-brain voxel-wise analyses: one parametric modulation analysis focussing on stimulus effects (i.e., the effect of reward/threat) and one general linear model (GLM) analysis focussing on response effects (i.e., the effect of the decision to approach or avoid). However, these additional analyses partially overlap with the BMM analyses, which, as explained above, are better suited for capturing higher-order interactions between task conditions and the observed behaviour. Therefore, we report the results of the BMM analyses. For completeness, voxel-wise model specifications and results are provided at: https://osf.io/hbjp5.

#### Subject-level analysis

After preprocessing, we employed subject-level models using SPM12 (Wellcome Trust Centre for Neuroimaging, London, UK). For each trial, we fitted one regressor for the *decision phase* (i.e., from trial onset until the response) and one regressor for the *outcome anticipation phase* (i.e., from response until the outcome), leading to 90 regressors for the *decision phase*, 90 regressors for the *outcome anticipation phase*. Additionally, we added regressors for each type of outcome (positive, negative, neutral) and motion regressors (3 translational, 3 rotational). This resulted in 3 additional regressors for the outcomes, 6 motion regressors, and 1 constant, leading to a total of 190 regressors. There was no censoring of data applied. The collinearity among all regressors of the *decision phase* and *outcome anticipation phase* was within an acceptable range (*M* = 0.0135, *Mdn* = 0.0056, *SD* = 0.0368, *min* = 0, *max* = 0.4025, see Mumford et al., 2015), with the highest collinearity observed between the regressors of the *decision phase* and *outcome anticipation phase* within the same trial. For each trial, we contrasted the regressor of each phase against the regressors of the corresponding phase in all other trials. Finally, trial-by-trial mean betas for each phase were extracted for each ROI using the MarsBaR toolbox (Brett et al., 2002). To ensure that the findings from our ROI analyses mainly stem from within-region voxels and not adjacent white matter or cerebrospinal fluid, we repeated our BMM analyses on the unsmoothed data from our ROI masks. These control analyses confirmed all key findings. Consequently, we report the results of the preregistered analyses on the smoothed data, as this approach accounts for anatomical variability and thereby yields more statistical power (Mikl et al., 2008).

#### Group-level analysis

For each phase, we employed a Bayesian Multivariate Multilevel (BMM) analysis where we used the trial-by-trial betas of our ROIs as dependent variables and reward, threat, decision, and their interactions as predictors. Given the correlational nature of our analysis, the choice whether variables are entered as dependent variables or predictors does not impact the results. However, since neural activity across different ROIs is correlated, using a multivariate model that accounts for this variance provides the best model fit and most accurate inferences. For full model specifications, see equation (2) below. Other model specifications were as described in the section ‘Bayesian multilevel analyses (general)’ above.

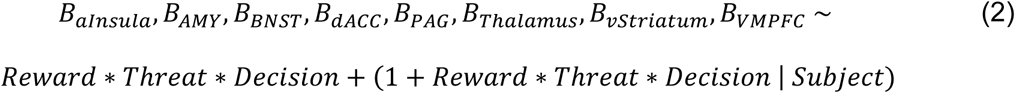

Given that the amygdala consists of functionally distinct anatomical subregions (McDonald, 1998; Price, 2003; Bzdok et al., 2013), we explored whether significant findings in the amygdala could be attributed to distinct contributions from its subnuclei: amygdalostriatal (AStr), basolateral (BLA), centromedial (CMA), and superficial (SFA) as defined in the SPM Anatomy Toolbox (Amunts et al., 2005; Eickhoff et al., 2007), see equation (3) below.

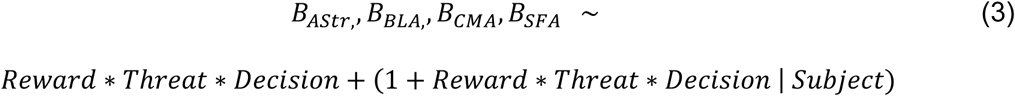

In the event of any significant interactions between predictors in our BMMs, planned comparisons were conducted using the emmeans package (Lenth et al., 2018) to clarify the nature of these interactions. Specifically, we investigated the interactions between reward and decision, as well as threat and decision, by examining whether the slope of reward was significant for each type of decision (approach/avoid) and whether the slope of threat was significant for each type of decision (approach/avoid). For interactions between reward, threat, and decision, we assessed whether the slope of threat was significantly different between approach and avoidance decisions in low and high reward conditions separately. If a significant difference was found within a specific reward level, we further examined whether the slope of threat was significant for each type of decision (approach/avoid) within that reward level. Betas and posterior credible intervals of all main effects, interaction effects, and post-hoc comparisons are provided at: https://osf.io/hbjp5.

## RESULTS

### Avoidance behaviour

As expected, participants avoided significantly more with decreasing reward levels (*B* =-2.01, 99.9% CI [−3.49, −0.74]) and increasing threat levels (*B* = 1.79, 99.9% CI [0.86, 2.75], see Figure 3). In low threat conditions, the effect of reward appears to be less pronounced, as participants typically approach, while in high reward conditions, participants tend to approach regardless of threat. However, this was not reflected in a significant interaction between reward and threat on avoidance (*B* = −.01, 90% CI [−0.19, 0.15]).

**Figure 3.**
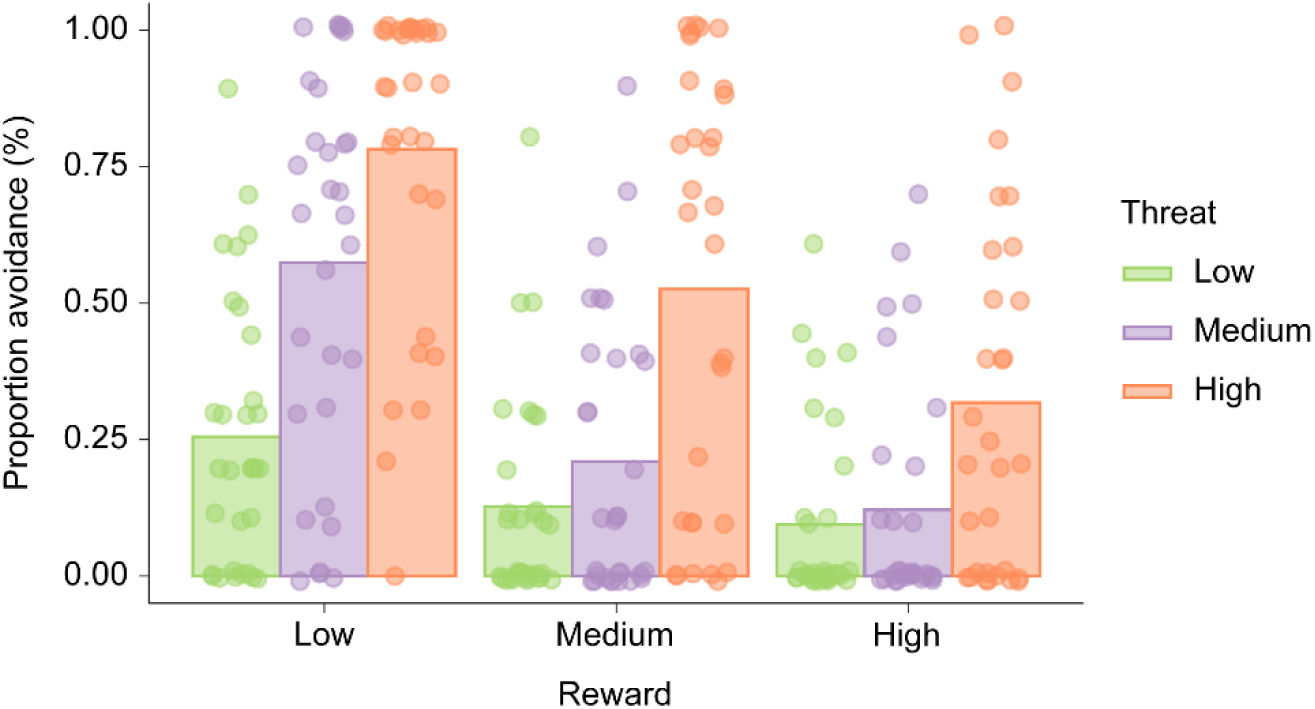
Overview task effects on avoidance behaviour. The bars represent the mean proportion avoidance as function of reward and threat level, overlaid with individual data points to illustrate variance between participants. Results showed the hypothesized opposing influences of reward and threat on avoidance.

### fMRI: decision phase

*Whole-brain voxel-wise analysis: decision phase vs baseline* Initial whole-brain voxel-wise analyses confirmed global activation across regions of interest hypothesized to be associated with weighing of reward and threat (aInsula, amygdala, BNST, dACC, PAG, thalamus, and vStriatum) when making approach-avoidance decisions (vs baseline; see Figure 4). Subsequently, to capture the weighing of rewards and threats for approach and avoidance decisions for each region of interest, we employed Bayesian Multivariate Multilevel (BMM) analyses.

**Figure 4.**
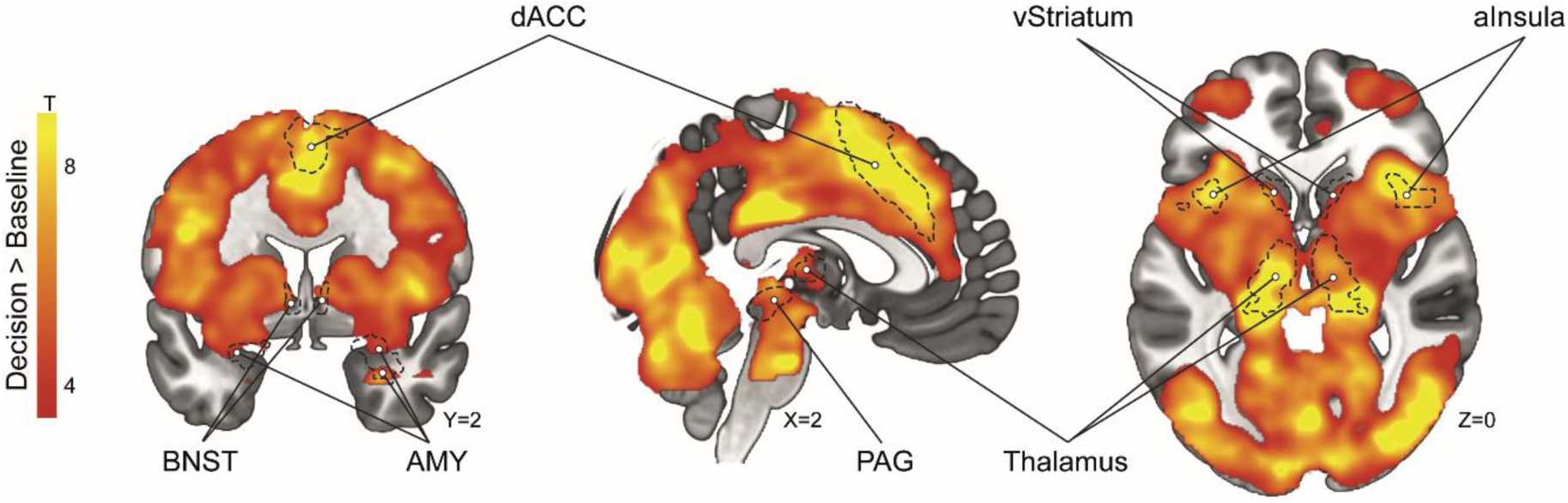
Whole-brain fMRI results demonstrating widespread activations during approach-avoidance decision-making (contrast: decision phase vs baseline). Images are shown at a cluster threshold of *p*<.05 corrected for multiple comparisons (FWE) after an initial threshold of *p*<.001 uncorrected. All marked regions of interests reach significance after correction for multiple comparisons (*p*FWE<.05). aInsula = anterior insula, AMY = amygdala, BNST = bed nucleus of the stria terminalis, dACC = dorsal anterior cingulate cortex, PAG = periaqueductal gray, vStriatum = ventral striatum.

### Bayesian Multivariate Multilevel fMRI analysis: reward, threat, and decision effects

First, we investigated the effects of reward, threat, decision, and their interactions on BOLD activity within our ROIs during the *decision phase*, i.e., prior to indicating approach-avoidance decisions (see Figure 5). We found that BOLD activity in the bed nucleus of the stria terminalis (BNST), periaqueductal gray (PAG), and ventral striatum increased as a function of reward, whereas BOLD activity in the dorsal anterior cingulate (dACC) increased as a function of threat. We did not find significant differences in overall BOLD activity between approach and avoidance decisions, nor significant interactions between reward and threat for any ROIs independent of decision (all 90% credible intervals overlapping with 0).

**Figure 5.**
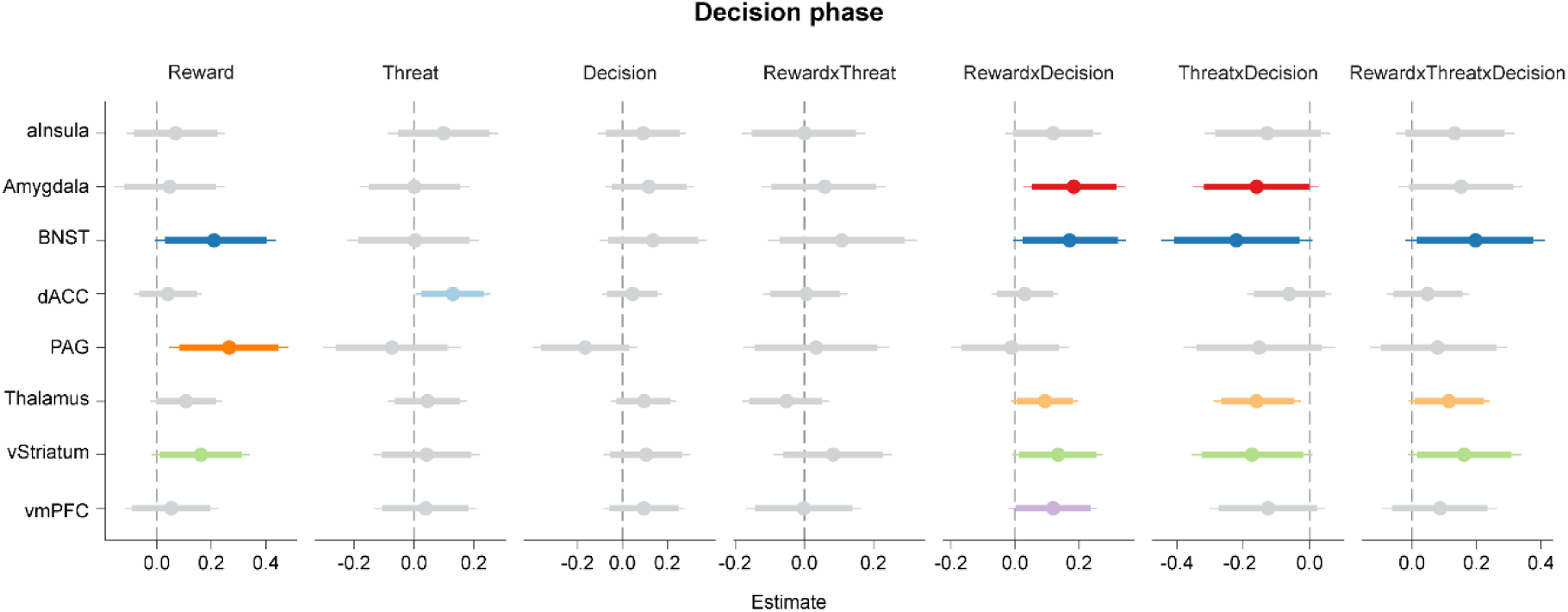
Posterior estimates and credible intervals for task-related activations in the regions of interest during the *decision phase*. This figure illustrates the posterior estimates (dots) with respect to the null (vertical dashed line). Bold lines represent 90% posterior credible intervals, while thin lines represent 95% posterior credible intervals. Significant effects, defined as ≥90% posterior credible intervals non-overlapping with zero, are highlighted in color. aInsula = anterior insula, BNST = bed nucleus of the stria terminalis, dACC = dorsal anterior cingulate cortex, PAG = periaqueductal gray, vmPFC = ventral medial prefrontal cortex, vStriatum = ventral striatum.

More importantly, we discovered that in several regions, the effect of reward and threat on BOLD activity differed between subsequent approach and avoidance decisions. Specifically, both the amygdala and vmPFC exhibited stronger BOLD activity with increasing reward preceding risky approach, but not risk-diminishing avoidance decisions (reward x decision; see Figure 6A), in line with the idea that activity in these regions may drive approach responses with increasing reward. Exploratory analyses indicated that this effect was present for all amygdala subnuclei (*B*_Astr_ = 0.05, 95% CI [0.00, 0.10]; *B*_BLA_ = 0.05, 95% CI [0.00, 0.10]; *B*_CMA_ = 0.05, 90% CI [0.00, 0.07]; *B*_SFA_ = 0.05, 90% CI [0.01, 0.09]). Moreover, stronger BOLD activity was observed in the amygdala in response to increasing threat for risk-diminishing avoidance relative to risky approach decisions (threat x decision; see Figure 6B), thus suggestive of a dual role for the amygdala in processing both rewards and threats. This effect was particularly driven by the amygdalostriatal subregion of the amygdala (*B*_Astr_ = −0.05, 95% CI [−0.09, −0.01], 90% credible intervals for all other subregions overlapping with 0). Interestingly, in addition to these two-way interactions, BOLD activity in the BNST, thalamus and ventral striatum exhibited a pattern suggesting that the integration of reward and threat differed for approach and avoidance decisions (reward x threat x decision; see Figure 6C). Specifically, when participants decided to avoid in low reward conditions, BOLD activity in the thalamus and ventral striatum was stronger with increasing threat. In contrast, prior to indicating risky approach decisions in low reward conditions, BOLD activity in the BNST and ventral striatum was decreased with increasing threat. Conversely, no differences in threat reactivity between approach and avoidance decisions were found when reward was high and the impact of threat on decisions much smaller (see Figure 3).

**Figure 6.**
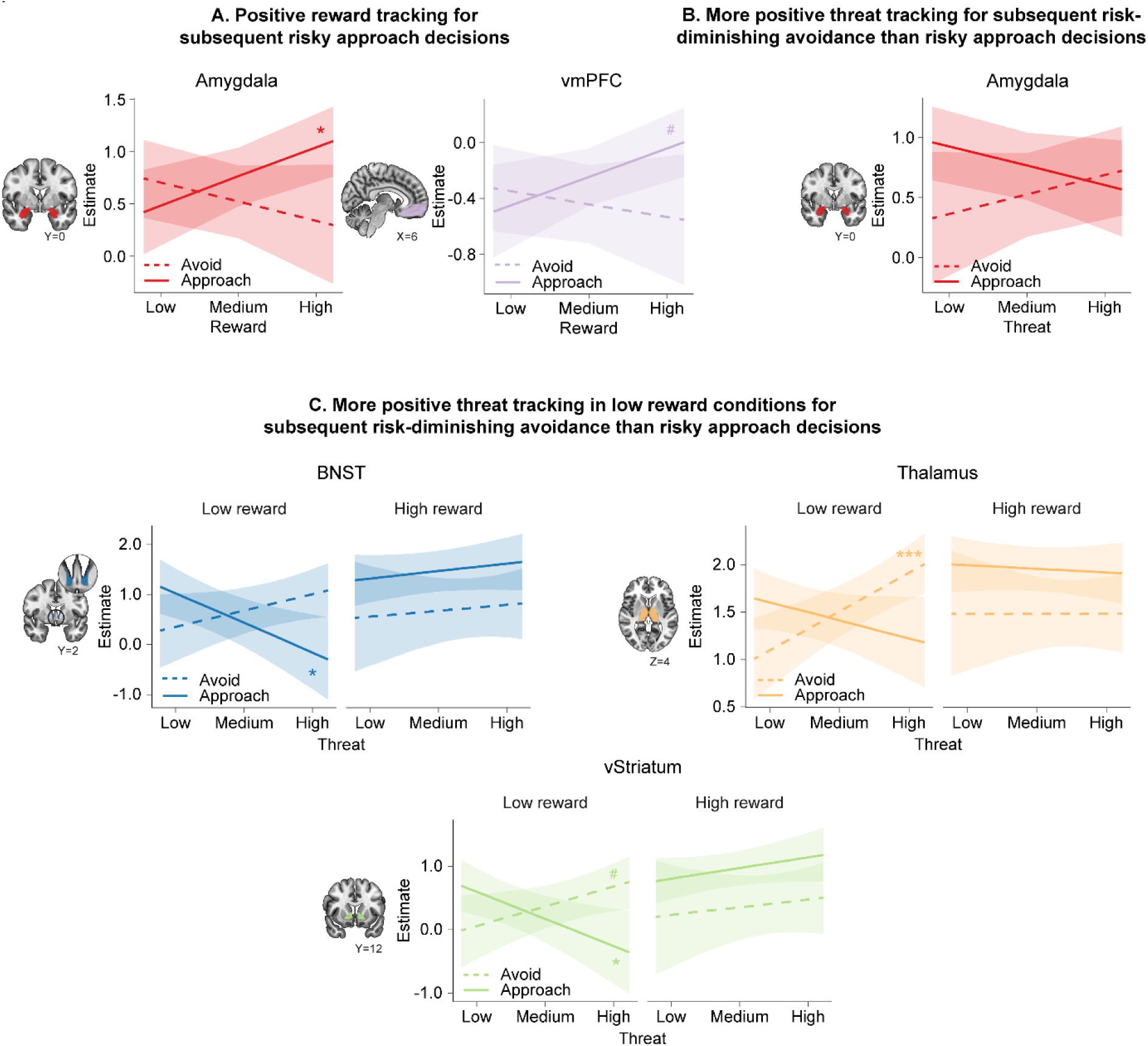
Effects of reward and threat on neural activity during approach-avoidance decision-making. The figure presents marginal effects plots for interactions between reward, threat, and decision in the *decision phase*. These plots depict separately for approach (solid lines) and avoidance (dashed lines): **(A)** the effect of *reward*, **(B)** the effect of *threat*, and **(C)** the *interaction between reward and threat* across all regions of interest where these interactions were observed. For each region of interest, only the highest-order interaction is plotted as it conveys most information. The asterisks indicate the significance levels of the slopes of **(A)** reward and **(B, C)** threat for approach and avoidance separately. #≥90%, *≥95%, **≥99%, or ***≥99.9% posterior credible intervals non-overlapping with zero. vmPFC = ventomedial prefrontal cortex, BNST = bed nucleus of the stria terminalis, vStriatum = ventral striatum.

In conclusion, our findings reveal that BOLD activity as a function of reward and threat preceding approach-avoidance decisions varied depending on the subsequent decision. Interestingly and contrary to theoretical predictions, several subcortical regions exhibited a pattern suggesting an integrative role.

### fMRI: outcome anticipation phase *Bayesian Multivariate Multilevel fMRI analysis: reward, threat, and decision effects*

Recent research suggests involvement of specific brain regions may depend on the specific stage of the approach-avoidance decision-making process. Therefore, we subsequently probed the contribution of our ROIs at the stage following approach-avoidance decisions, when participants were passively anticipating the outcome. As indicated above, risky approach decisions were associated with a relatively high chance of receiving either electric stimulation or monetary reward (40% for each; high risk/high gain), whereas risk-diminishing avoidance decisions were associated with a relatively low chance of receiving these outcomes (10% for each; low risk/low gain). During this *outcome anticipation phase*, BOLD activity in the amygdala, BNST, PAG, and thalamus was more pronounced following risky approach decisions compared to risk-diminishing avoidance decisions (see Figure 7). Moreover, we observed stronger BOLD activity in the aInsula and PAG with increasing threat. We did not find any significant main effects of reward nor interactions between reward and threat in any ROI (all 90% credible intervals overlapping with 0).

**Figure 7.**
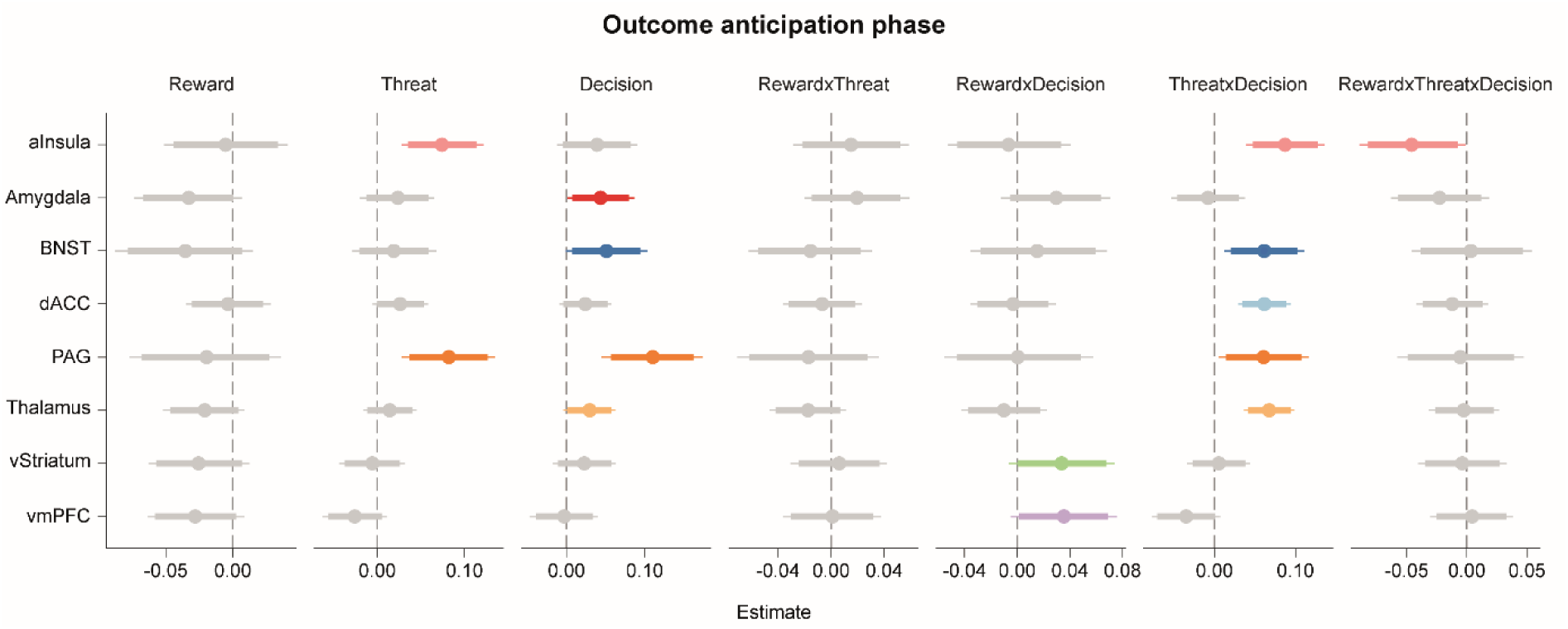
Posterior estimates and credible intervals for task-related activations in the regions of interest during the *outcome anticipation phase*. This figure illustrates the posterior estimates (dots) with respect to the null (vertical dashed line). Bold lines represent 90% posterior credible intervals, while thin lines represent 95% posterior credible intervals. Significant effects, defined as ≥90% posterior credible intervals non-overlapping with zero, are highlighted in color. aInsula = anterior insula, BNST = bed nucleus of the stria terminalis, dACC = dorsal anterior cingulate cortex, PAG = periaqueductal gray, vmPFC = ventral medial prefrontal cortex, vStriatum = ventral striatum.

Interestingly, in several regions, the influence of reward and threat on the BOLD response during the outcome anticipation phase depended on the preceding decision. Following risk-diminishing avoidance decisions, when the chance of receiving shocks or rewards was low, BOLD activity within the vStriatum and vmPFC significantly decreased with increasing reward. Conversely, no such effects were observed following risky approach decisions, when the chance of receiving shocks or rewards was relatively high (reward x decision; see Figure 8A). Furthermore, we found that following risk-diminishing avoidance decisions, BOLD activity in the thalamus decreased with increasing threat. In contrast, following risky approach decisions, when the risk of receiving electrical stimulation was high, BOLD activity in the aInsula, BNST, dACC, PAG, and thalamus increased with increasing threat (threat x decision; see Figure 8B). This finding further aligns with the presumed role of these regions in anticipating impending threats. Notably, there was integration of reward and threat in the aInsula, exhibiting variations between approach and avoidance decisions (reward x threat x decision; see Figure 8C). This interaction was driven by the fact that while anticipating the outcome of avoidance decisions (low risk), BOLD activity in the aInsula was marginally decreased as a function of threat in low reward conditions. In contrast, while anticipating the outcome of approach decisions (high risk), BOLD activity in the aInsula increased as a function of threat in low reward conditions. Thus, the pattern of BOLD activity in the aInsula was similar to patterns observed in other threat anticipation regions. However, it was the only region that exclusively exhibited this pattern when reward was low.

**Figure 8.**
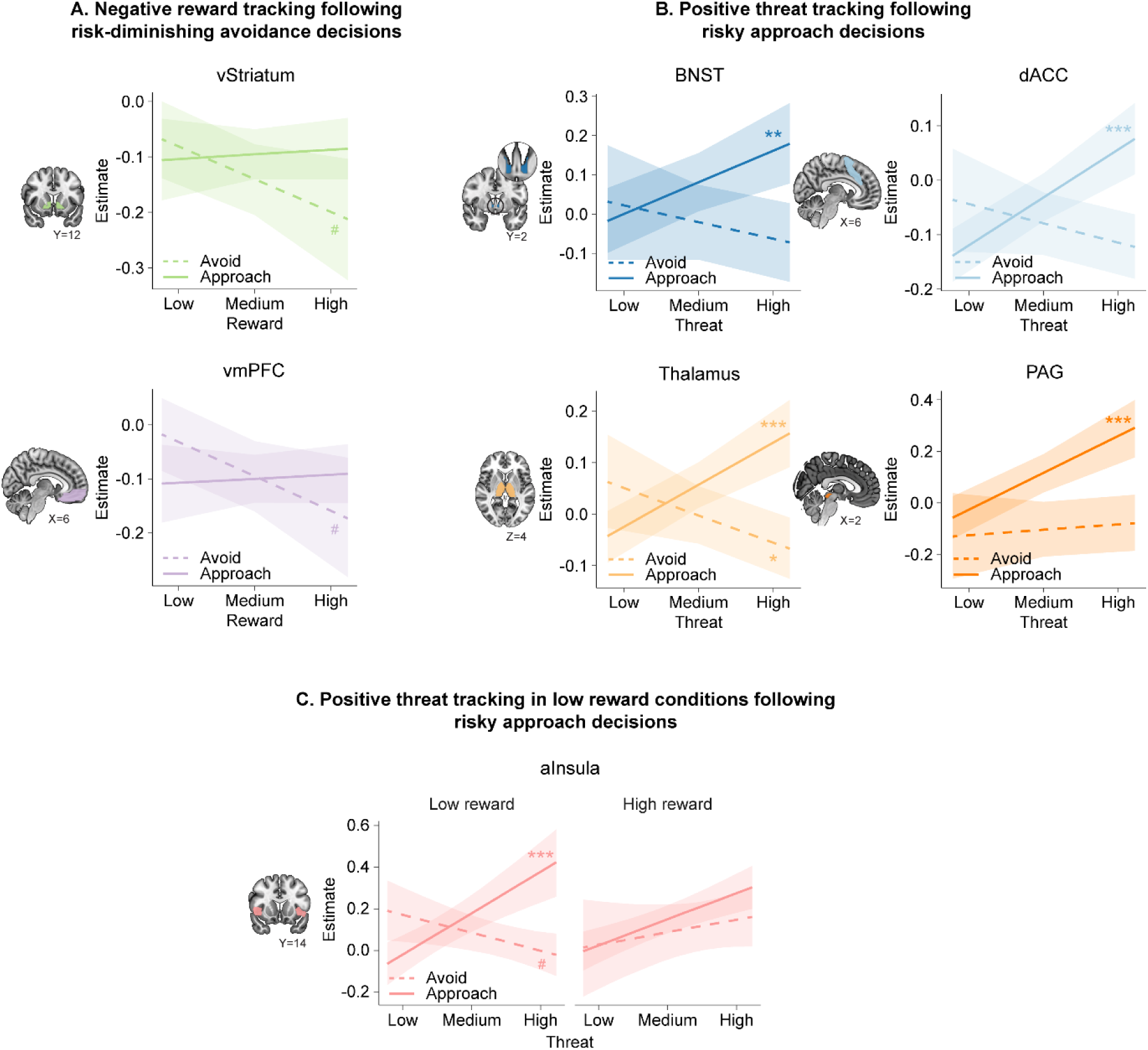
Effects of reward and threat on neural activity during anticipation of the outcome of approach-avoidance decisions. The figure presents marginal effects plots for interactions between reward, threat, and decision in the *outcome anticipation phase*. These plots depict separately for approach (solid lines) and avoidance (dashed lines): **(A)** the effect of *reward*, **(B)** the effect of *threat*, and **(C)** the *interaction between reward and threat* across all regions of interest where these interactions were observed. For each region of interest, only the highest-order interaction is plotted as it conveys most information. The asterisks indicate the significance levels of the slopes of **(A)** reward and **(B, C)** threat for approach and avoidance separately. #≥90%, *≥95%, **≥99%, or ***≥99.9% posterior credible intervals non-overlapping with zero. vStriatum = ventral striatum, vmPFC = ventromedial prefrontal cortex, BNST = bed nucleus of the stria terminalis, dACC = dorsal anterior cingulate cortex, PAG = periaqueductal gray, aInsula = anterior insula.

In conclusion, following avoidance decisions, when the chance of receiving rewards was relatively low, regions within the reward network negatively tracked reward magnitude. Conversely, after approach decisions, when the risk of receiving electric shock was relatively high, regions in the salience network positively tracked threat magnitude, with the aInsula only doing so when reward was low.

## DISCUSSION

In this study, we investigated how the decision to engage in risky approach or risk-diminishing avoidance may arise from distributed neural processing of reward and threat information. We aimed to provide a better understanding of the neural dynamics governing approach-avoidance decisions by 1) investigating whether the decision to approach or avoid is preceded by parallel and/or integrated neural processing of reward and threat across brain regions previously implicated in approach-avoidance decision-making and 2) evaluated how such processing and integration evolve over time following approach-avoidance decisions during outcome anticipation. Our preregistered Bayesian Multivariate Multilevel (BMM) analyses on fMRI data revealed two key findings. First, in contrast to the notion of *parallel processing* of reward and threat in subcortical regions traditionally associated predominantly with either reward (vStriatum, vmPFC) or threat (amygdala, BNST) processing, we found evidence for *dual* and *integrated processing*. Specifically, there was *dual processing* of reward and threat information in the amygdala and evidence for *integration* in other subcortical regions, including the vStriatum, thalamus, and BNST. Critically, the weighing of reward and threat in these regions varied as a function of the subsequent decision to approach or avoid. Second, after indicating the decision to approach or avoid, we observed that neural activity in regions traditionally associated with reward (vmPFC, vStriatum) and threat (BNST) processing, as well as the salience network (dACC, thalamus, PAG), was influenced the magnitudes of reward and threat, consistent with previous theories. Together, these findings illuminate neural dynamics of approach-avoidance decision-making and suggest distributed subcortical integration of reward and threat as a potential driver of subsequent approach-avoidance decisions.

In accordance with prior research, we found that when making decisions involving risky approach or risk-dimishing avoidance, individuals carefully balance potential rewards and threats (Talmi et al., 2009; Basten et al., 2010; Park et al., 2011; Aupperle et al., 2015; Schlund et al., 2016; Hulsman et al., 2021a, 2024; Klaassen et al., 2021; Moughrabi et al., 2022). To delve deeper into the distributed neural patterns underlying this decision-making process, we employed Bayesian Multivariate Multilevel (BMM) analyses. The observed neural patterns deviated from conventional perspectives that primarily suggest isolated reward and threat processing in subcortical regions, followed by the integration of reward and threat in cortical regions such as the ACC or vmPFC (Yacubian et al., 2006; Talmi et al., 2009; Basten et al., 2010; Park et al., 2011; Azab and Hayden, 2020; Livermore et al., 2021). Instead, our findings suggest that the decision to approach or avoid is supported by partially overlapping subcortical regions sensitive to both reward and threat information. Specifically, the amygdalostriatal subnucleus of the amygdala, BNST, thalamus, and vStriatum showed a pattern of increasing neural activity as function of threat prior to indicating risk-diminishing avoidance as compared to risky approach decisions. In contrast, the vmPFC uniquely displayed a reward-by-decision effect without being moderated by threat, consistent with its hypothesized role in reward valuation (Talmi et al., 2009; Aupperle and Paulus, 2010; Basten et al., 2010; Bartra et al., 2013) and subsequent approach decisions (Pedersen et al., 2021). The convergence of brain regions engaged in processing both reward and threat corresponds with recent findings demonstrating that the BNST, vStriatum, and PAG respond to both reward and threat information (Murty et al., 2023) and that conflict between approach and avoidance tendencies is represented at the level of the vStriatum (Ironside et al., 2020). Additionally, converging findings have shown that the amygdala responds to both aversive and appetitive stimuli (Baxter and Murray, 2002; O’Doherty, 2004; Schultz, 2006; Pessoa, 2010).

Our findings, which highlight subcortical integration of reward and threat, provide a new perspective that contrasts with prior human neuroimaging research. However, they do resonate with long-standing reports in rodents (Gentry et al., 2016; Burgos-Robles et al., 2017; Hamel et al., 2017; Beyeler et al., 2018), collectively suggesting that the traditional notion of these regions responding primarily to either reward or threat is untenable. Nevertheless, it remains possible that depending on the context certain regions predominantly encode reward or threat information, as observed here, with vmPFC and PAG demonstrating sensitivity to reward and the dACC to threat. Tentatively, such signals could serve as input for regions integrating both reward and threat inputs, such as the BNST, vStriatum, and thalamus. This perspective aligns with theories proposing that these regions, due to their connectivity profile, may serve as intermediaries between lower downstream subcortical and upstream cortical regions (Avery et al., 2014; Goode and Maren, 2017; De Groote and de Kerchove d’Exaerde, 2021; Hammack et al., 2021; Sieveritz and Raghavan, 2021), akin to striatal-thalamo-cortical loops in motor control (Pessoa, 2023).

Furthermore, the observed increase in threat-related activity in the vStriatum, thalamus, and BNST preceding avoidance relative to approach under low reward, aligns with recent findings that increased threat representations in these regions during decision-making were linked to avoidance (Moughrabi et al., 2022). Behaviourally, the presence of high reward attenuated the impact of threat on subsequent decisions. This attenuation was also reflected in neural responses within the vStriatum, thalamus, and BNST. These findings further align with previous research outside the field of value-based decision making that demonstrate competition between reward and threat processing in various brain regions, including the BNST (Choi et al., 2014).

Finally, we demonstrated shifts in these neural patterns of reward/threat processing following approach-avoidance decisions. Following risky approach decisions, we observed positive tracking of threat in the salience and threat network, encompassing the BNST, thalamus, dACC, and PAG, all of which have consistently been associated with threat anticipation (Mechias et al., 2010; Fullana et al., 2016; Klumpers et al., 2017; Andrzejewski et al., 2019; Patrick et al., 2019). Interestingly, in the aInsula we only found increased activity as a function of threat following approach decisions when rewards were low. This aligns with previous research suggesting that the presence of reward can suppress the effect of threat in the aInsula (Cristofori et al., 2015). Conversely, following risk-diminishing avoidance decisions we found negative tracking of reward in the reward network, including the vStriatum and vmPFC. These findings appear to reflect decreased reward expectancies after avoidance decisions (Heekeren et al., 2007; Jia et al., 2016; Pujara et al., 2016; Rehbein et al., 2023).

Despite the implementation of a fast multiband-multiecho fMRI sequence (MB3ME3), further delineation the temporal dynamics of approach-avoidance decisions would require methods with superior temporal resolution, such as MEG or EEG (Khemka et al., 2017; McFadyen et al., 2023). Another limitation of our study is a relatively small sample size (N=28 included in MRI analyses). Our analyses were preregistered, and we employed BMM analyses to investigate neural responses to minimize the risk of false positives (Kajimura et al., 2023). Still, replication of our findings is desirable, particularly in clinical populations that show approach-avoidance deficits to gain further knowledge of how reward-threat balances across distinct regions may be disturbed (Ironside et al., 2020; Smith et al., 2021b, 2021a, 2023; McDermott et al., 2022).

In conclusion, our findings unveiled distributed cortico-subcortical processing and subcortical integration of reward and threat prior to the decision to approach or avoid. These findings suggest a departure from traditional, yet widely professed, views that segregate brain regions as either predominantly reward-sensitive or threat-sensitive, and assign the integration of reward and threat primarily to cortical regions. Following approach-avoidance decisions, however, brain regions traditionally associated with reward and threat processing reflected reward and threat tracking contingent on the decision that was made. These findings provide new theoretical insight into the unfolding neural dynamics of approach-avoidance decision-making.

## Supporting information

Supplemental information and results

