## Supplemental information and results for "How distributed subcortical integration of reward and threat may inform subsequent approach-avoidance decisions"

**Supplement**

**Atypical response patterns**

As preregistered (Hulsman et al., 2021b) atypical response patterns yielded: 50% more avoidance responses were made in low threat conditions than high threat condition (i.e., low reward/low threat > low reward/high threat, medium reward/low threat > medium reward/high threat, high reward/low threat > high reward/high threat) and 50% or more avoidance decisions were made in high reward condition than low reward conditions (i.e., high reward/low threat > low reward/low threat, high reward/medium threat > low reward/medium threat, high reward/high threat > low reward/high threat). These criteria are in line with our previous studies (Hulsman et al., 2021a, 2024).

**Whole-brain fMRI analysis: general task effects**

To investigate general task effects, we ran a parametric modulation analysis. Predictors of this analysis were the *decision phase* (i.e., from trial onset until the response) and the *outcome anticipation phase* (i.e., from response until the outcome). For these two predictors reward level, threat level, and the interaction between reward and threat were included as linear parametric modulation regressors. Other predictors were the outcomes: positive, negative, and neutral. As recommended by Mumford et al. (2015), all parametric regressors were orthogonalized with respect to the unmodulated trial regressor per subject (i.e., demeaned). Realignment parameters were included to reduce movement-related artefacts. High-pass filtering at 1/128 Hz and a first-order autoregressive model were used as standard. General task effects independent of reward, threat, and decision were investigated by contrasting neural activity during the *decision phase* against baseline and contrasting neural activity during the *outcome anticipation phase* against baseline. The baseline consisted of neural responses during the inter-trial interval (ITI). Finally, responses during the outcome delivery were investigated by contrasting positive against negative outcomes. All single-subject contrast maps were subsequently subjected to a t-test. All fMRI figures were created in MRIcroGL, with use of the MNI 152 template image. As preregistered, whole-brain results were familywise (FWE) corrected for multiple comparisons according to random field theory ($p<.05$). Small volume FWE correction ($p<.05$) was applied to all ROIs after an initial threshold of $p<.005$ uncorrected. In the event of significant amygdala activations, we used the SPM anatomy toolbox to determine which particular amygdala subregion was activated. This toolbox provides a likelihood index ($P_{excess}$) indicating the probability of activation in a particular amygdala subregion. $P_{excess}$ is based on the degree of overlap between the observed activation and the probability maps of the centromedial amygdala (CMA), superficial amygdala

(SFA), and basolateral amygdala (BLA). $P_{excess}$ > 1 indicates a high probability that the observed activation belongs to that particular subregion, while $P_{excess}$ < 1 indicates that the observed activation primarily intersects with peripheral regions (Amunts et al., 2005; Eickhoff et al., 2007). Note that this analysis is exploratory as we had no a priori hypotheses regarding the specific roles of individual amygdala subregions.

**Whole brain fMRI analysis: stimulus and response focused models**

As mentioned in the manuscript, due to unbalanced data, conventional MRI analyses are unable to simultaneously model both stimulus and response effects. Therefore, we conducted two separate voxel-wise analyses one focussing on stimulus effects (i.e., the effect of reward/threat) and one on response effects (i.e., the effect of the decision to approach or avoid). For both models realignment parameters were included to reduce movement-related artefacts. High-pass filtering at $^{1}/_{128}$ Hz and a first-order autoregressive model were used as standard. All fMRI figures were created in MRIcroGL, with use of the MNI 152 template image. As preregistered, whole-brain results were familywise (FWE) corrected for multiple comparisons according to random field theory ($p$<.05) after an initial threshold of $p$<.001 uncorrected. Small volume FWE correction ($p$<.05) was applied to all ROIs after an initial threshold of $p$<.005 uncorrected.

*Stimulus focused model*

To investigate the effects of reward and threat on BOLD signal variation, we ran a parametric modulation analysis. This parametric modulation analysis is described above (see p1). For both phases (*decision phase* and *outcome anticipation phase*), we performed a one-sample t-test on the first-level univariate parametric modulation beta maps.

*Response focused model*

A general linear model was composed to relate BOLD signal variation to the decision that participants made (approach/avoid). Predictors of this model were approach and avoid. These predictors were modelled for two phases (*decision* and *outcome anticipation*) separately. Other predictors were the outcomes: positive, negative, neutral. The effect of decision was investigated using a t-test on contrasts comparing approach vs avoidance decisions during each of these phases.

### Results

#### *fMRI: decision phase*

*Whole-brain voxelwise analysis: general task effects (decision phase vs baseline) independent of reward, threat, and decision*

Whole-brain analyses revealed that prior to indicating approach-avoidance decisions, there was increased neural activity in a broad network of regions, including, but not limited to regions typically associated with threat processing (amygdala, particularly in the right BLA: $P_{excess}$ = 1.45) and the salience network (anterior insula, dACC, thalamus) (see Table S1). Region of interest analyses further revealed significant activations in the BNST, PAG and vStriatum (see Table S2 for ROI results).

**Table S1. Whole-brain results for BOLD responses during the *decision phase***

| | Peak coordinates, MNI | | | | | |
|---|---|---|---|---|---|---|
| | x | y | z | $k_E$ | Z | $p_{FWE\text{-}corr}$ |
| ***Decision phase > Baseline*** | | | | | | |
| **L+R dACC, L+R thalamus,** L postcentral gyrus, L+R middle occipital gyrus, L+R cerebellum, R fusiform gyrus, L+R lingual gyrus, L precentral gyrus, L inferior parietal gyrus, L superior parietal gyrus, L fusiform gyrus, L calcarine, L+R SMA | 4 | -32 | 30 | 402450 | 7.77 | <.001 |
| **R anterior insula**, R middle frontal gyrus, R insula, R inferior frontal gyrus (opercular), R precentral gyrus, R inferior frontal gyrus (triangular part), R putamen, R rolandic operculum, R inferior frontal gyrus (pars orbitalis), R superior frontal gyrus | 42 | 32 | 26 | 3423 | 7.62 | <.001 |
| **L anterior insula**, L precentral gyrus, L putamen, L rolandic operculum, L inferior frontal gyrus (pars orbitalis), L pallidum, L middle frontal gyrus, L superior temporal pole, L inferior frontal gyrus (triangular part) | -50 | 4 | 28 | 2307 | 7.07 | <.001 |
| R parahippocampal gyrus, R fusiform gyrus, **R amygdala** | 30 | -2 | -32 | 55 | 6.24 | <.001 |
| R supramarginal gyrus, R temporal superior gyrus | 56 | -42 | 32 | 227 | 6.02 | <.001 |
| L middle frontal gyrus, L inferior frontal gyrus (triangular), L inferior frontal gyrus (opercular) | -46 | 40 | 22 | 893 | 5.88 | <.001 |
| L fusiform gyrus | -32 | -4 | -30 | 8 | 5.38 | .005 |
| R putamen, R pallidum | 22 | -4 | 8 | 45 | 5.34 | <.001 |
| R precentral gyrus, R postcentral gyrus | 22 | -32 | 70 | 18 | 5.31 | .001 |
| R superior frontal gyrus | 18 | 24 | 58 | 5 | 5.30 | .010 |
| R middle cingulum, R anterior cingulum | 4 | 0 | 30 | 9 | 5.29 | .004 |
| R caudate | 10 | 4 | 6 | 12 | 5.04 | .003 |
| L insula | -38 | 4 | -12 | 2 | 4.93 | .021 |
| R superior frontal gyrus | 30 | 52 | 12 | 1 | 4.92 | .029 |
| L inferior frontal gyrus (triangular part) | -48 | 48 | 8 | 4 | 4.88 | .012 |
| R middle frontal gyrus | 30 | 62 | 4 | 1 | 4.85 | .029 |
| L middle cingulum | -14 | -34 | 50 | 1 | 4.84 | .029 |

All results are whole-brain voxelwise FWE-corrected p<.05 and described in order of strongest activation (Z-values). Regions of interest are highlighted in bold font. L=left, R=right

**Table S2. ROI results for BOLD responses for general task effects during the *decision phase***

| | Peak coordinates, MNI | | | | | |
|---|---|---|---|---|---|---|
| | x | y | z | $k_E$ | Z | $p_{FWE-corr}$ |
| ***Decision phase > Baseline*** | | | | | | |
| L+R thalamus | -20 | -30 | 2 | 2201 | 7.68 | <.001 |
| L+R dACC | 2 | 6 | 54 | 2831 | 7.42 | <.001 |
| R aInsula | 34 | 18 | 2 | 319 | 7.20 | <.001 |
| L aInsula | -36 | 12 | 6 | 302 | 6.65 | <.001 |
| R amygdala | 32 | 0 | -30 | 166 | 5.50 | <.001 |
| L vStriatum | -16 | 10 | -4 | 190 | 5.30 | <.001 |
| R BNST | 8 | 4 | 6 | 11 | 5.03 | <.001 |
| L+R PAG | 4 | -26 | -6 | 55 | 4.96 | <.001 |
| L BNST | -8 | 4 | 6 | 18 | 4.95 | <.001 |
| L amygdala | -22 | 0 | -12 | 128 | 4.71 | <.001 |
| R vStriatum | 18 | 14 | -2 | 136 | 4.26 | .002 |
| L amygdala | -30 | -2 | -28 | 2 | 3.05 | .075* |

All results are FWE small volume-corrected p<.05 for bilateral anatomical mask after an initial voxel wise threshold of p<.005. Results are described in order of strongest activation (Z values). L=left, R=right. *Marginally significant (between p=.05 and .10).

*Stimulus-related effects (parametric modulation of reward and threat)*

Whole-brain parametric modulation analyses revealed that during the decision phase, BOLD signal in several brain regions, including, but not limited to, regions typically associated with the reward network (bilateral ventral striatum), the salience network (bilateral (anterior) insula, bilateral mid/anterior cingulum, bilateral midbrain), bilateral supplementary motor area, bilateral precuneus, and bilateral cerebellum, were positively correlated with the reward level of that trial. Interestingly, similar reward effects were observed in regions typically associated with the threat network (right amygdala, bilateral BNST). Contrary to our expectations, we did not find significant correlations with threat. See Table S3 for whole-brain results and S4 for ROI results.

**Table S3. Whole-brain results for BOLD responses associated with reward and threat during the *decision phase***

| | Peak coordinates, MNI | | | | | |
|---|---|---|---|---|---|---|
| | x | y | z | $k_E$ | Z | $p_{FWE-corr}$ |
| ***Decision phase – reward correlation (positive):*** | | | | | | |
| L+R postcentral gyrus/L+R rolandic operculum/L+R precuneus/L+R precentral gyrus/L+R supplementary motor area/L+R superior temporal gyrus/L middle cingulum | 66 | -2 | 14 | 20915 | 5.16 | <.001 |
| R temporal pole (superior temporal gyrus)/R amygdala/R parahippocampal gyrus/R fusiform gyrus/R hippocampus/R temporal pole (middle temporal gyrus)/R inferior frontal gyrus (pars orbitalis) | 32 | 14 | -28 | 215 | 4.23 | .096 |
| **L+R ventral striatum**/L+R caudate/L+R olfactory cortex/L putamen/L pallidum/R rectus/L hipopocampus | 6 | 8 | -6 | 452 | 4.07 | .006 |
| L middle frontal gyrus/L superior frontal gyrus | -32 | 34 | 34 | 223 | 3.95 | .087 |
| L cuneus/L superior occipital gyrus/L superior parietal gyrus/L precuneus/L middle occipital gyrus | -14 | -76 | 38 | 575 | 3.85 | .002 |
| L fusiform gyrus/L+R cerebellum/L inferior temporal gyrus | -38 | -34 | -32 | 234 | 3.71 | .075 |

All results are whole-brain voxelwise FWE-corrected p<.05 and described in order of strongest activation (Z values). L=left, R=right. Non-significant effects *decision phase*: reward correlation negative, threat correlation positive, threat correlation negative.

**Table S4. Region of interest results for BOLD responses of parametric modulation maps reward and threat during the *decision phase***

| | Peak coordinates, MNI | | | | | |
|---|---|---|---|---|---|---|
| | x | y | z | $k_E$ | Z | $p_{FWE-corr}$ |
| ***Decision phase - reward correlation (positive)*** | | | | | | |
| L thalamus | -20 | -34 | 4 | 433 | 4.42 | .002 |
| L+R dACC | 0 | 4 | 50 | 272 | 4.09 | .009 |
| R vStriatum | 6 | 8 | -6 | 152 | 4.07 | .003 |
| R amygdala | 32 | -2 | -28 | 145 | 3.95 | .003 |
| R thalamus | 16 | -32 | -2 | 58 | 3.95 | .012 |
| L vStriatum | -8 | 6 | -2 | 141 | 3.85 | .006 |
| L BNST | -8 | 4 | -2 | 16 | 3.66 | .003 |
| R vmPFC | 10 | 14 | -14 | 37 | 3.63 | .056* |
| R PAG | 4 | -30 | -8 | 9 | 3.58 | .003 |
| R BNST | 6 | 4 | -2 | 20 | 3.39 | .006 |
| R aINS | 44 | 10 | 0 | 59 | 3.20 | .045 |
| L aINS | -46 | 8 | -4 | 75 | 3.18 | .049 |
| L amygdala | -20 | -6 | -12 | 30 | 3.07 | .054* |
| L PAG | -2 | -28 | -6 | 7 | 3.00 | .015 |

All results are FWE small volume-corrected p<.05 for bilateral anatomical mask after an initial voxel wise threshold of p<.005. Results are described in order of strongest activation (Z values). L=left, R=right. *Marginally significant (between p=.05 and .10). Non-significant effects *decision phase*: reward correlation negative, threat correlation negative, threat correlation positive.

*Response-related BOLD signal variation (approach vs avoid)*

Whole-brain analyses revealed that prior to indicating the decision to approach or avoid, BOLD signal in the executive-control network (comprising the left inferior/middle/superior frontal gyrus, as well as the left precentral gyrus and left superior parietal gyrus) was decreased for avoidance decisions compared to approach decisions. No significant clusters were identified where the BOLD signal increased for avoidance decisions in comparison to approach decisions. See Table S5 for whole-brain results and S6 for ROI results.

**Table S5. Whole-brain results for BOLD responses associated with approach-avoidance behaviour during the *decision phase***

| | Peak coordinates, MNI | | | | | |
|---|---|---|---|---|---|---|
| | x | y | z | $k_E$ | Z | $p_{FWE\text{-}corr}$ |
| ***Decision phase – approach > avoid*** | | | | | | |
| L middle frontal gyrus/L precentral gyrus/L superior frontal gyrus | -36 | 8 | 58 | 475 | 4.33 | .003 |
| L inferior parietal gyrus/L angular gyrus/L superior parietal gyrus/L superior occipital gyrus | -50 | -52 | 52 | 833 | 4.18 | <.001 |
| L inferior frontal gyrus (triangular part)/L middle frontal gyrus | -44 | 44 | 18 | 294 | 4.11 | .027 |

All results are whole-brain voxelwise FWE-corrected p<.05 and described in order of strongest activation (Z values). L=left, R=right. Non-significant effects *decision phase*: avoid > approach.

**Table S6. Region of interest results for BOLD responses associated with approach-avoidance behaviour during the *decision phase***

| | Peak coordinates, MNI | | | | | |
|---|---|---|---|---|---|---|
| | x | y | z | $k_E$ | Z | $p_{FWE\text{-}corr}$ |
| ***Decision phase – avoid < approach*** | | | | | | |
| L thalamus | -18 | -34 | 2 | 35 | 3.94 | .013 |
| R vmPFC | 18 | 14 | -20 | 32 | 3.72 | .046 |
| R vStriatum | 8 | 14 | 4 | 9 | 3.00 | .087* |

All results are FWE small volume-corrected p<.05 for bilateral anatomical mask after an initial voxel wise threshold of p<.005. Results are described in order of strongest activation (Z values). L=left, R=right.* Marginally significant (between p=.05 and .10). *Marginally significant (between p=.05 and .10). Non-significant effects *decision phase*: avoid > approach.

### Outcome anticipation phase

*Whole-brain voxelwise analysis: general task effects (outcome anticipation phase vs baseline) independent of reward, threat, and decision*

Whole-brain analyses showed that while anticipating the outcome of approach-avoidance decisions, there was increased neural activity in regions that are part of the limbic system (hippocampus, thalamus) compared to baseline (see Figure S1 and Table S7). Other regions that showed with similar activation patterns were located in the occipital and temporal lobes, as well as the cerebellum. In addition, region of interest analyses revealed increased activity in the anterior insula during the outcome anticipation phase (see Table S8 for ROI results).

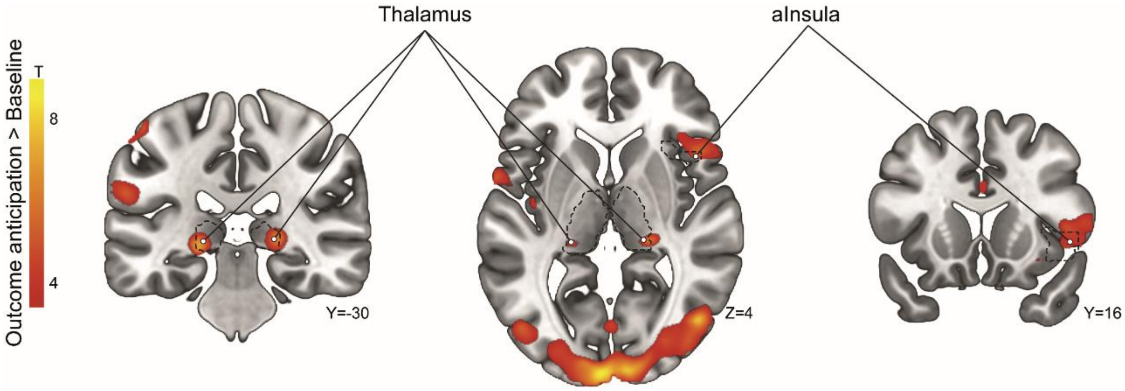


**Figure S1. Whole-brain fMRI results for the contrast outcome anticipation vs baseline.** For illustrative purposes, images are shown at a threshold of $p<.001$ uncorrected. All marked regions of interests reach significance after correction for multiple comparisons ($p_{FWE}<.05$).

**Table S7. Whole-brain results for BOLD responses during the *outcome anticipation phase***

| | Peak coordinates, MNI | | | | | |
|---|---|---|---|---|---|---|
| | x | y | z | $k_E$ | Z | $p_{FWE-corr}$ |
| ***Outcome anticipation phase > Baseline*** | | | | | | |
| R inferior occipital gyrus, L+R calcarine, R fusiform gyrus, R lingual gyrus, R cuneus, L+R superior occipital gyrus, L+R middle occipital gyrus, R inferior temporal gyrus, R middle temporal gyrus, L+R cerebellum | 38 | -76 | -8 | 1650 | 6.26 | <.001 |
| L fusiform gyrus, L lingual gyrus, L cerebellum | -38 | -82 | -18 | 142 | 5.55 | <.001 |
| L hippocampus, **L thalamus** | -20 | -30 | -2 | 15 | 5.42 | .003 |
| R inferior frontal gyrus (triangular part), R inferior frontal gyrus (opercular part) | 46 | 20 | 8 | 8 | 5.22 | .007 |
| L supramarginal gyrus | -62 | -22 | 28 | 6 | 5.05 | .010 |
| **R thalamus** | 22 | -30 | 2 | 8 | 5.04 | .007 |
| R middle occipital gyrus | 30 | -92 | 20 | 1 | 4.79 | .031 |

All results are whole-brain voxelwise FWE-corrected p<.05 and described in order of strongest activation (Z-values). Regions of interest are highlighted in bold font. L=left, R=right.

**Table S8. ROI results for BOLD responses for general task effects during the *outcome anticipation phase***

| | Peak coordinates, MNI | | | | | |
|---|---|---|---|---|---|---|
| | x | y | z | $k_E$ | Z | $p_{FWE-corr}$ |
| ***Outcome anticipation > Baseline*** | | | | | | |
| L thalamus | -20 | -30 | -2 | 69 | 5.42 | <.001 |
| R thalamus | 22 | -30 | 2 | 70 | 5.04 | <.001 |
| R aInsula | 48 | 16 | 4 | 102 | 4.04 | .004 |
| L amygdala | -18 | 0 | -20 | 17 | 3.10 | .060* |

All results are FWE small volume-corrected p<.05 for bilateral anatomical mask after an initial voxel wise threshold of p<.005. Results are described in order of strongest activation (Z values). L=left, R=right.* Marginally significant (between p=.05 and .10)

#### *Stimulus-related effects (parametric modulation of reward and threat)*

Whole-brain parametric modulation analyses showed that after indicating the decision to approach or avoid, there was a negative correlation between reward level and BOLD signal across a wide range of brain regions, such as the left hippocampus, the bilateral occipital lobe, bilateral cerebellum, left temporal lobe, left supplementary motor area, left ventromedial prefrontal cortex, right insula, and right thalamus.

In addition, we found a positive correlation between threat and BOLD signal in the bilateral cerebellum, salience network (bilateral anterior cingulate cortex, bilateral anterior insula, right thalamus), bilateral midbrain, and bilateral basal ganglia. Unexpectedly, on whole-brain level we found no evidence for amygdala or BNST activation as function of threat. Nevertheless, ROI analyses revealed a significant threat effect in the right amygdala. See Table S9 for whole-brain results and S10 for ROI results.

**Table S9. Whole-brain results for BOLD responses associated with reward and threat during the *outcome anticipation phase***

| | Peak coordinates, MNI | | | | | |
|---|---|---|---|---|---|---|
| | x | y | z | $k_E$ | Z | $p_{FWE\text{-}corr}$ |
| ***Outcome anticipation phase – reward correlation (negative):*** | | | | | | |
| L hippocampus/L parahippocampal gyrus/L fusiform gyrus/L cerebellum | -24 | -14 | -18 | 480 | 5.16 | .001 |
| R lingual gyrus/L+R calcarine/L+R cerebellum/L+R precuneus/R fusiform gyrus/R cuneus/L+R vermis/L+R posterior cingulate gyrus | 14 | -88 | 2 | 3027 | 5.09 | <.001 |
| L middle temporal gyrus/L postcentral gyrus/L inferior temporal gyrus/L superior temporal gyrus/L cerebellum/L fusiform gyrus | -66 | -24 | -2 | 1488 | 4.71 | <.001 |
| L+R postcentral gyrus/R precentral gyrus/L+R paracentral lobule/R superior temporal gyrus/R rolandic operculum/R inferior parietal gyrus/R supplementary motor area/L+R precuneus/R middle temporal gyrus | -6 | -34 | 56 | 3670 | 4.68 | <.001 |
| L lingual gyrus/L precuneus/L calcarine/L cerebellum/L fusiform gyrus/L hippocampus/L parahippocampal gyrus | -14 | -46 | -2 | 439 | 4.64 | .001 |
| L cerebellum | -30 | -70 | -42 | 449 | 4.63 | .001 |
| L middle occipital gyrus/L superior occipital gyrus/L cuneus/L calcarine | -22 | -82 | 22 | 924 | 4.56 | <.001 |
| R middle occipital gyrus/R superior parietal gyrus/R superior occipital gyrus/R angular gyrus/R cuneus/R precuneus/R middle temporal gyrus | 22 | -80 | 48 | 957 | 4.50 | <.001 |
| R middle frontal gyrus/R superior frontal gyrus | 28 | 22 | 56 | 588 | 4.45 | <.001 |
| L inferior frontal gyrus (triangular part)/L inferior frontal gyrus (pars orbitalis)/L inferior frontal gyrus (opercular part) | -48 | 42 | -4 | 518 | 4.41 | <.001 |
| L middle frontal gyrus/L precentral gyrus/L superior frontal gyrus | -30 | 14 | 56 | 458 | 4.21 | .001 |
| R insula/R putamen/R Heschl's gyrus/R rolandic operculum/R superior temporal gyrus/R pallidum | 38 | -22 | 6 | 384 | 4.17 | .003 |
| **L ventromedial prefrontal cortex** | -14 | 46 | -22 | 272 | 4.04 | .014 |
| ***Outcome anticipation – threat correlation (positive)*** | | | | | | |
| L+R cerebellum/L+R vermis/R lingual gyrus/R fusiform gyrus | 28 | -56 | -24 | 2537 | 5.32 | <.001 |
| **R insula**/R pallidum/R putamen/R inferior frontal gyrus (pars orbitalis)/R inferior frontal gyrus (opercular part)/R temporal pole (superior temporal gyrus) | 36 | 8 | -4 | 1129 | 5.27 | <.001 |
| L postcentral gyrus/L precentral gyrus/L paracentral lobule/L supplementary motor area/L+R mid cingulum/L+R anterior cingulum/L superior parietal gyrus/L+R superior frontal gyrus | -38 | -34 | 72 | 4549 | 4.94 | <.001 |
| L cerebellum | -34 | -50 | -32 | 689 | 4.85 | <.001 |
| **L insula**/L putamen/L pallidum/L rolandic operculum/L superior temporal gyrus/L temporal pole (superior temporal gyrus)/L inferior frontal gyrus (triangular part)/L inferior frontal gyrus (pars orbicularis) | -34 | 12 | 6 | 1308 | 4.82 | <.001 |
| L supramarginal gyrus/L superior temporal gyrus/L rolandic operculum/L postcentral gyrus/L middle temporal gyrus | -50 | -26 | 22 | 595 | 4.36 | <.001 |
| L+R brain stem | -6 | -24 | -16 | 558 | 4.26 | <.001 |
| R middle temporal gyrus/R superior temporal gyrus | 46 | -30 | -2 | 155 | 4.24 | .080* |
| L middle frontal gyrus/L superior frontal gyrus | -30 | 44 | 22 | 199 | 4.56 | .033 |
| L calcarine/L lingual gyrus | -4 | -86 | 0 | 184 | 3.87 | .044 |

All results are whole-brain voxelwise FWE-corrected p<.05 and described in order of strongest activation (Z values). L=left, R=right. Non-significant effects *outcome anticipation phase*: reward correlation positive, threat correlation negative.

**Table S10. Region of interest results for BOLD responses of parametric modulation maps reward and threat during the *outcome anticipation phase***

| | Peak coordinates, MNI | | | | | |
|---|---|---|---|---|---|---|
| | x | y | z | $k_E$ | Z | $p_{FWE\text{-}corr}$ |
| ***Outcome anticipation phase – reward correlation (positive)*** | | | | | | |
| R dACC | 12 | 10 | 68 | 13 | 3.57 | .067* |
| ***Outcome anticipation phase – reward correlation (negative)*** | | | | | | |
| R thalamus | 16 | -24 | 6 | 120 | 4.24 | .006 |
| L vmPFC | -14 | 46 | -22 | 638 | 4.04 | .021 |
| L vStriatum | -6 | 12 | -8 | 35 | 3.55 | 0.22 |
| R amygdala | 22 | -4 | -18 | 44 | 3.46 | .022 |
| R vStriatum | 10 | 10 | -12 | 3 | 3.11 | .078* |
| L amygdala | -20 | -8 | -16 | 8 | 2.98 | .084* |
| ***Outcome anticipation phase – threat correlation (positive)*** | | | | | | |
| R aINS | 38 | 10 | -4 | 286 | 5.09 | <.001 |
| L aINS | -36 | 14 | 4 | 252 | 4.73 | <.001 |
| L+R dACC | 8 | 20 | 30 | 871 | 4.53 | .003 |
| L thalamus | -14 | -18 | -2 | 153 | 4.07 | .012 |
| R thalamus | 8 | -8 | -4 | 251 | 3.90 | .021 |
| L PAG | -2 | -26 | -4 | 30 | 3.83 | .002 |
| R amygdala | 32 | 4 | -20 | 2 | 2.96 | .095* |
| ***Outcome anticipation phase – threat correlation (negative)*** | | | | | | |
| L+R vmPFC | -2 | 38 | -14 | 256 | 3.95 | .032 |

All results are FWE small volume-corrected p<.05 for bilateral anatomical mask after an initial voxel wise threshold of p<.005. Results are described in order of strongest activation (Z values). L=left, R=right. *Marginally significant (between p=.05 and .10).

#### *Response-related effects (approach vs avoid)*

After making approach-avoidance decisions, whole-brain analyses revealed that BOLD signal in a distributed network of brain regions, encompassing frontal, parietal, temporal, and cingulate (bilateral middle/anterior cingulate cortex) regions was increased for avoidance decisions compared to avoidance decisions. No significant clusters were identified where the BOLD signal decreased for avoidance decisions compared to approach decisions. See Table S11 for whole-brain results and S12 for ROI results.

**Table 11. Whole-brain results for BOLD responses associated with approach-avoidance behaviour during the *outcome anticipation phase***

| | Peak coordinates, MNI | | | | | |
|---|---|---|---|---|---|---|
| | x | y | z | $k_E$ | Z | $p_{FWE\text{-}corr}$ |
| ***Outcome anticipation phase – avoid > approach*** | | | | | | |
| R angular gyrus/R middle occipital gyrus/R inferior parietal gyrus/R superior occipital gyrus/R superior parietal gyrus | 56 | -64 | 36 | 1316 | 5.08 | <.001 |
| R middle frontal gyrus/R middle frontal gyrus (pars orbitalis)/R superior frontal gyrus/R inferior frontal gyrus (pars orbitalis)/R inferior frontal gyrus (triangular part) | 32 | 62 | 4 | 700 | 4.74 | <.001 |
| R middle frontal gyrus/R superior frontal gyrus/R medial superior frontal gyrus/R inferior frontal gyrus (triangular part)/R supplementary motor area | 14 | 38 | 56 | 1649 | 4.74 | <.001 |
| L angular gyrus/L inferior parietal gyrus/L supramarginal gyrus | -48 | -60 | 46 | 863 | 4.73 | <.001 |
| R inferior temporal gyrus/R middle temporal gyrus | 68 | -46 | -16 | 713 | 4.71 | <.001 |
| L+R middle cingulum/L post cingulum/L precuneus | 2 | -32 | 44 | 251 | 4.45 | .019 |
| L middle temporal gyrus/L inferior temporal gyrus | -60 | -26 | -8 | 520 | 4.33 | <.001 |
| L middle frontal gyrus/L precentral gyrus/L superior frontal gyrus | -38 | 20 | 54 | 756 | 4.25 | <.001 |
| L superior frontal gyrus (medial)/L+R middle cingulum/**L anterior cingulum** | -2 | 34 | 38 | 191 | 4.18 | .055 |
| L middle frontal gyrus/L superior frontal gyrus | -34 | 58 | 8 | 295 | 4.11 | .009 |
| L inferior frontal gyrus (pars orbitalis)/L middle frontal gyrus (pars orbitalis) | -48 | 42 | -6 | 188 | 3.62 | .058 |

All results are whole-brain voxelwise FWE-corrected p<.05 and described in order of strongest activation (Z values). L=left, R=right. Non-significant effects *outcome anticipation phase*: approach > avoid.

**Table S12. Region of interest results for BOLD responses associated with approach-avoidance behaviour during the *outcome anticipation phase***

| | Peak coordinates, MNI | | | | | |
|---|---|---|---|---|---|---|
| | x | y | z | $k_E$ | Z | $p_{FWE\text{-}corr}$ |
| ***Outcome anticipation phase – avoid > approach*** | | | | | | |
| L+R dACC | -2 | 34 | 38 | 192 | 5.01 | .009 |
| R thalamus | 22 | -22 | 6 | 54 | 4.28 | .037 |
| L thalamus | -24 | -32 | 6 | 19 | 4.11 | .052* |
| R dACC | 20 | 14 | 58 | 18 | 4.03 | .074* |
| L aINS | -48 | 18 | -4 | 10 | 3.32 | .096* |
| R aINS | 50 | 18 | -6 | 4 | 3.31 | .095* |

All results are FWE small volume-corrected p<.05 for bilateral anatomical mask after an initial voxel wise threshold of p<.005. Results are described in order of strongest activation (Z values). L=left, R=right. *Marginally significant (between p=.05 and .10). *Marginally significant (between p=.05 and .10). Non-significant effects *outcome anticipation phase*: avoid < approach.

### Outcome delivery

#### *Negative > positive*

At whole-brain level, receiving negative outcomes, as opposed to positive outcomes, was associated with increased neural activity in various brain regions distributed across the brain. These regions consisted of the insula, auditory-motor regions (Heschl's gyrus, superior temporal gyrus), sensorimotor regions (pre- and postcentral gyrus), supplementary motor area, cerebellum, rolandic operculum, and supramarginal gyrus. Additionally, region of interest analyses revealed similar activation patterns in the amygdala, dACC, thalamus, and PAG (see Table S13 for whole-brain results and S14 for ROI results).

*Positive > negative*

Receiving positive outcomes, as opposed to negative outcomes, was associated with increased neural responses in the default mode network, including the precuneus, hippocampus, and parahippocampal gyrus. In addition, the middle temporal gyrus and fusiform gyrus showed similar increases in neural activity for receiving positive compared to negative outcomes. Furthermore, region of interest analyses showed increased activity in the vmPFC for positive compared to negative outcomes (see Table S13 for whole-brain results and S14 for ROI results).

**Table S13. Whole-brain results for BOLD responses during *outcome delivery***

| | Peak coordinates, MNI | | | | | |
|---|---|---|---|---|---|---|
| | x | y | z | $k_E$ | Z | $p_{FWE\text{-}corr}$ |
| ***Outcome: negative > positive*** | | | | | | |
| **R (anterior) insula**, R superior temporal gyrus, R rolandic operculum, R putamen, R Heschl's gyrus | 40 | -8 | -4 | 733 | 7.28 | <.001 |
| **L (anterior) insula**, L rolandic operculum, L supramarginal gyrus, L superior temporal gyrus, L Heschl's gyrus, L postcentral gyrus | -40 | -6 | -6 | 1661 | 7.12 | <.001 |
| R supramarginal gyrus, R rolandic operculum, R superior temporal gyrus | 58 | -24 | 24 | 248 | 5.89 | <.001 |
| L postcentral gyrus | -38 | -28 | 50 | 33 | 5.57 | <.001 |
| R cerebellum | 20 | -54 | -18 | 29 | 5.20 | <.001 |
| L precentral gyrus, L postcentral gyrus | -44 | -24 | 62 | 13 | 5.09 | .003 |
| L postcentral gyrus | -48 | -30 | 60 | 6 | 4.95 | .010 |
| L precentral gyrus | -42 | -18 | 66 | 3 | 4.94 | .018 |
| R cerebellum | 14 | -64 | -42 | 2 | 4.93 | .023 |
| L middle cingulum | -2 | 10 | 38 | 6 | 4.91 | .010 |
| L supplemental motor area | -12 | 0 | 74 | 2 | 4.88 | .023 |
| L postcentral gyrus | -22 | -40 | 76 | 1 | 4.85 | .031 |
| L tectum midbrain | -6 | -30 | -6 | 1 | 4.81 | .031 |
| L superior parietal gyrus | -20 | -44 | 74 | 2 | 4.81 | .023 |
| ***Outcome positive > negative*** | | | | | | |
| R fusiform gyrus, R parahippocampal gyrus | 30 | -30 | -20 | 46 | 5.40 | <.001 |
| L precuneus | -4 | -58 | 36 | 12 | 5.15 | .004 |
| L parahippocampal gyrus, L hippocampus | -20 | -16 | -22 | 8 | 5.14 | .007 |
| L precuneus | -2 | -54 | 28 | 2 | 4.88 | .023 |
| L middle temporal gyrus | -64 | -14 | -16 | 1 | 4.81 | .031 |

All results are whole-brain voxelwise FWE-corrected p<.05 and described in order of strongest activation (Z values). Regions of interest are highlighted in bold font. L=left, R=right.

**Table S14 ROI results for BOLD responses during *outcome delivery***

| | Peak coordinates, MNI | | | | | |
|---|---|---|---|---|---|---|
| | x | y | z | $k_E$ | Z | $p_{FWE\text{-}corr}$ |
| ***Negative > Positive*** | | | | | | |
| R aInsula | 38 | 12 | 2 | 233 | 5.84 | <.001 |
| L aInsula | -36 | 8 | 0 | 213 | 5.15 | <.001 |
| L dACC | -2 | 10 | 40 | 250 | 4.76 | .001 |
| R dACC | 12 | 12 | 58 | 292 | 4.01 | .016 |
| L thalamus | -8 | -6 | 8 | 328 | 3.84 | .023 |
| L amygdala | -22 | 0 | -16 | 67 | 3.78 | .008 |
| L dACC | -8 | 4 | 72 | 36 | 3.62 | .056* |
| R amygdala | 20 | 4 | -16 | 29 | 3.34 | .031 |
| R amygdala | 34 | 4 | -20 | 4 | 3.16 | .051* |
| L+R PAG | -2 | -32 | -6 | 9 | 2.97 | .019 |
| ***Positive > Negative*** | | | | | | |
| L+R vmPFC | -10 | 34 | -12 | 473 | 4.63 | .002 |

All results are FWE small volume-corrected p<.05 for bilateral anatomical mask after an initial voxel wise threshold of p<.005. Results are described in order of strongest activation (Z values). L=left, R=right. *Marginally significant (between p=.05 and .10).

**Table S15. Model results *decision phase***

| ROI | Reward | Threat | Decision | Reward x Threat | Reward x Decision | Threat x Decision | Reward x Threat x Decision |
|---|---|---|---|---|---|---|---|
| aINS | *B*=0.07<br>$CI_{90\%}$ = [-0.08, 0.22] | *B*=0.10<br>$CI_{90\%}$ = [-0.05, 0.25] | *B*=0.09<br>$CI_{90\%}$ = [-0.07, 0.26] | *B*=0.00<br>$CI_{90\%}$ = [-0.15, 0.15] | *B*=0.15<br>$CI_{90\%}$ = [-0.01, 0.30] | *B*=-0.13<br>$CI_{90\%}$ = [-0.29, 0.03] | *B*=0.13<br>$CI_{90\%}$ = [-0.02, 0.29] |
| Amygdala | *B*=0.05<br>$CI_{90\%}$ = [-0.12, 0.22] | *B*=0.00<br>$CI_{90\%}$ = [-0.15, 0.16] | *B*=0.12<br>$CI_{90\%}$ = [-0.05, 0.29] | *B*=0.06<br>$CI_{90\%}$ = [-0.09, 0.21] | ***B*=0.23**<br>**$CI_{95\%}$ = [0.03, 0.43]** | ***B*=-0.16**<br>**$CI_{90\%}$ = [-0.32, -0.00]** | *B*=0.15<br>$CI_{90\%}$ = [-0.01, 0.31] |
| BNST | ***B*=0.21**<br>**$CI_{90\%}$ = [0.03, 0.40]** | *B*=0.00<br>$CI_{90\%}$ = [-0.19, 0.18] | *B*=0.14<br>$CI_{90\%}$ = [-0.06, 0.34] | *B*=0.11<br>$CI_{90\%}$ = [-0.07, 0.29] | ***B*=0.21**<br>**$CI_{90\%}$ = [0.03, 0.40]** | ***B*=-0.22**<br>**$CI_{90\%}$ = [-0.41, -0.03]** | ***B*=0.20**<br>**$CI_{90\%}$ = [0.01, 0.38]** |
| dACC | *B*=0.04<br>$CI_{90\%}$ = [-0.06, 0.15] | ***B*=0.13**<br>**$CI_{95\%}$ = [0.01, 0.25]** | *B*=0.05<br>$CI_{90\%}$ = [-0.07, 0.16] | *B*=0.00<br>$CI_{90\%}$ = [-0.10, 0.10] | *B*=0.04<br>$CI_{90\%}$ = [-0.07, 0.15] | *B*=-0.06<br>$CI_{90\%}$ = [-0.17, 0.05] | *B*=0.05<br>$CI_{90\%}$ = [-0.06, 0.16] |
| PAG | ***B*=0.26**<br>**$CI_{95\%}$ = [0.05, 0.48]** | *B*=-0.07<br>$CI_{90\%}$ = [-0.26, 0.11] | *B*=-0.17<br>$CI_{90\%}$ = [-0.36, 0.03] | *B*=0.03<br>$CI_{90\%}$ = [-0.14, 0.22] | *B*=-0.02<br>$CI_{90\%}$ = [-0.21, 0.17] | *B*=-0.15<br>$CI_{90\%}$ = [-0.34, 0.04] | *B*=0.08<br>$CI_{90\%}$ = [-0.10, 0.26] |
| Thalamus | *B*=0.11<br>$CI_{90\%}$ = [-0.00, 0.22] | *B*=0.05<br>$CI_{90\%}$ = [-0.06, 0.15] | *B*=0.09<br>$CI_{90\%}$ = [-0.03, 0.22] | *B*=-0.05<br>$CI_{90\%}$ = [-0.16, 0.06] | ***B*=0.12**<br>**$CI_{90\%}$ = [0.01, 0.23]** | ***B*=-0.16**<br>**$CI_{95\%}$ = [-0.29, -0.03]** | ***B*=0.11**<br>**$CI_{90\%}$ = [0.01, 0.22]** |
| vStriatum | ***B*=0.16**<br>**$CI_{90\%}$ = [0.01, 0.31]** | *B*=0.04<br>$CI_{90\%}$ = [-0.11, 0.19] | *B*=0.11<br>$CI_{90\%}$ = [-0.05, 0.27] | *B*=0.08<br>$CI_{90\%}$ = [-0.06, 0.23] | ***B*=0.17**<br>**$CI_{90\%}$ = [0.02, 0.32]** | ***B*=-0.17**<br>**$CI_{90\%}$ = [-0.33, -0.02]** | ***B*=0.16**<br>**$CI_{90\%}$ = [0.02, 0.31]** |
| vmPFC | *B*=0.05<br>$CI_{90\%}$ = [-0.09, 0.20] | *B*=0.04<br>$CI_{90\%}$ = [-0.11, 0.18] | *B*=0.10<br>$CI_{90\%}$ = [-0.06, 0.25] | *B*=-0.00<br>$CI_{90\%}$ = [-0.14, 0.14] | ***B*=0.15**<br>**$CI_{90\%}$ = [0.00, 0.29]** | *B*=-0.13<br>$CI_{90\%}$ = [-0.27, 0.02] | *B*=0.09<br>$CI_{90\%}$ = [-0.06, 0.23] |

aINS = anterior insula, BNST = bed nucleus of the stria terminalis, dACC = dorsal anterior cingulate cortex, PAG = periaqueductal gray, vStriatum = ventral striatum, vmPFC = ventromedial prefrontal cortex. Effects that reached (marginal) significance (i.e., ≥90% posterior credible interval was non-overlapping with zero) are displayed in bold.

**Table S16. Post-hoc comparisons for the interaction between reward, threat, and decision: differences in threat effect between approach and avoid decisions in the *decision phase*, separated for low and high reward conditions.**

| | *Differences in threat effect between approach and avoidance decisions* | |
|---|---|---|
| | Low reward | High reward |
| BNST | ***B*=-0.91**<br>**$CI_{99\%}$ = [-1.76, -0.05]** | *B*=0.04<br>$CI_{90\%}$ = [-0.58, 0.64] |
| Thalamus | ***B*=-0.59**<br>**$CI_{99\%}$ = [-1.09, -0.11]** | *B*=-0.05<br>$CI_{90\%}$ = [-0.41, 0.31] |
| vStriatum | ***B*=-0.73**<br>**$CI_{99\%}$ = [-1.44, -0.08]** | *B*=0.04<br>$CI_{90\%}$ = [-0.44, 0.54] |

BNST = bed nucleus of the stria terminalis, vStriatum = ventral striatum. Effects that reached (marginal) significance (i.e., ≥90% posterior credible interval was non-overlapping with zero) are displayed in bold.

**Table S17. Post-hoc comparisons for the interaction between reward, threat, and decision: threat effect in low reward conditions in the *decision phase*, separated by decision (approach vs avoid).**

| | *Threat effect in low reward conditions* | |
|---|---|---|
| | Approach | Avoid |
| BNST | ***B*=-0.59**<br>**$CI_{95\%}$ = [-1.04, -0.10]** | *B*=0.33<br>$CI_{90\%}$ = [-0.04, 0.68] |
| Thalamus | *B*=-0.19<br>$CI_{90\%}$ = [-0.42, 0.04] | ***B*=0.41**<br>**$CI_{99.9\%}$ = [0.01, 0.86]** |
| vStriatum | ***B*=-0.42**<br>**$CI_{95\%}$ = [-0.79, 0.02]** | ***B*=0.31**<br>**$CI_{90\%}$ = [0.02, 0.60]** |

BNST = bed nucleus of the stria terminalis, vStriatum = ventral striatum. Effects that reached (marginal) significance (i.e., ≥90% posterior credible interval was non-overlapping with zero) are displayed in bold.

**Table S18. Post-hoc comparisons for the interaction between reward and decision: reward effect in the *decision phase*, separated by decision (approach vs avoid).**

| | *Reward effect* | |
|---|---|---|
| | Approach | Avoid |
| Amygdala | ***B*=0.28**<br>**$CI_{95\%}$ =[0.03, 0.51]** | *B*=-0.18<br>$CI_{90\%}$ = [-0.44, 0.10] |
| BNST | ***B*=0.42**<br>**$CI_{99\%}$ = [0.06, 0.76]** | *B*=-0.00<br>$CI_{90\%}$ = [-0.31, 0.29] |
| Thalamus | ***B*=0.22**<br>**$CI_{99\%}$ = [0.02, 0.42]** | *B*=-0.01<br>$CI_{90\%}$ = [-0.18, 0.18] |
| vStriatum | ***B*=0.33**<br>**$CI_{99\%}$ = [0.07, 0.62]** | *B*=-0.01<br>$CI_{90\%}$ = [-0.25, 0.23] |
| vmPFC | ***B*=0.20**<br>**$CI_{90\%}$ = [0.03, 0.37]** | *B*=-0.10<br>$CI_{90\%}$ = [-0.34, 0.13] |

BNST = bed nucleus of the stria terminalis, vStriatum = ventral striatum, vmPFC = ventromedial prefrontal cortex. Effects that reached (marginal) significance (i.e., ≥90% posterior credible interval was non-overlapping with zero) are displayed in bold.

**Table S19. Post-hoc comparisons for the interaction between threat and decision: threat effect in the *decision phase*, separated by decision (approach vs avoid).**

| | *Threat effect* | |
|---|---|---|
| | Approach | Avoid |
| Amygdala | *B*=-0.16<br>$CI_{90\%}$ = [-0.34, 0.02] | *B*=0.16<br>$CI_{90\%}$ = [-0.09, 0.41] |
| BNST | ***B*=-0.22**<br>**$CI_{90\%}$ = [-0.44, -0.00]** | *B*=0.22<br>$CI_{90\%}$ = [-0.07, 0.53] |
| Thalamus | *B*=-0.11<br>$CI_{90\%}$ = [-0.24, 0.01] | ***B*=0.21**<br>**$CI_{90\%}$ = [0.02, 0.38]** |
| vStriatum | *B*=-0.13<br>$CI_{90\%}$ = [-0.31, 0.05] | *B*=0.22<br>$CI_{90\%}$ =[-0.03, 0.46] |

BNST = bed nucleus of the stria terminalis, vStriatum = ventral striatum. Effects that reached (marginal) significance (i.e., ≥90% posterior credible interval was non-overlapping with zero) are displayed in bold.

**Table S20. Model results *outcome anticipation phase*.**

| ROI | Reward | Threat | Decision | Reward x threat | Reward x decision | Threat x decision | Reward x threat x decision |
|---|---|---|---|---|---|---|---|
| aInsula | *B*=-0.01 $CI_{90\%}$ = [-0.04, 0.03] | ***B*=0.07 $CI_{99.9\%}$= [0.00, 0.16]** | *B*=0.04 $CI_{90\%}$ = [-0.00, 0.08] | *B*=0.02 $CI_{90\%}$ = [-0.02, 0.05] | *B*=-0.01 $CI_{90\%}$ = [-0.05, 0.03] | ***B*=0.09 $CI_{99.9\%}$ = [0.00, 0.17]** | ***B*=-0.05 $CI_{95\%}$ = [-0.09, -0.00]** |
| Amygdala | *B*=-0.03 $CI_{90\%}$ = [-0.07, 0.00] | *B*=0.02 $CI_{90\%}$ = [-0.01, 0.06] | ***B*=0.04 $CI_{95\%}$ = [0.00, 0.09]** | *B*=0.02 $CI_{90\%}$ = [-0.01, 0.05] | *B*=0.03 $CI_{90\%}$ = [-0.01, 0.06] | *B*=-0.01 $CI_{90\%}$ = [-0.05, 0.03] | *B*=-0.02 $CI_{90\%}$ = [-0.06, 0.01] |
| BNST | *B*=-0.04 $CI_{90\%}$ = [-0.08, 0.01] | *B*=0.02 $CI_{90\%}$ = [-0.02, 0.06] | ***B*=0.05 $CI_{90\%}$ = [0.01, 0.09]** | *B*=-0.02 $CI_{90\%}$ = [-0.05, 0.02] | *B*=0.02 $CI_{90\%}$ = [-0.03, 0.06] | ***B*=0.06 $CI_{95\%}$ = [0.01, 0.12]** | *B*=0.00 $CI_{90\%}$ = [-0.04, 0.05] |
| dACC | *B*=-0.00 $CI_{90\%}$ = [-0.03, 0.02] | *B*=0.03 $CI_{90\%}$ = [-0.00, 0.05] | *B*=0.02 $CI_{90\%}$ = [-0.00, 0.05] | *B*=-0.01 $CI_{90\%}$ = [-0.03, 0.02] | *B*=-0.00 $CI_{90\%}$ = [-0.03, 0.02] | ***B*=0.06 $CI_{95\%}$ = [0.03, 0.09]** | *B*=-0.01 $CI_{90\%}$ = [-0.04, 0.01] |
| PAG | *B*=-0.02 $CI_{90\%}$ = [-0.07, 0.03] | ***B*=0.08 $CI_{99\%}$ = [0.01, 0.15]** | ***B*=0.11 $CI_{99.9\%}$ = [0.00, 0.22]** | *B*=-0.02 $CI_{90\%}$ = [-0.06, 0.03] | *B*=0.00 $CI_{90\%}$ = [-0.05, 0.05] | ***B*=0.06 $CI_{95\%}$ = [0.01, 0.12]** | *B*=-0.01 $CI_{90\%}$ = [-0.05, 0.04] |
| Thalamus | *B*=-0.02 $CI_{90\%}$ = [-0.05, 0.00] | *B*=0.01 $CI_{90\%}$ = [-0.01, 0.04] | ***B*=0.03 $CI_{90\%}$ = [0.00, 0.06]** | *B*=-0.02 $CI_{90\%}$ = [-0.04, 0.01] | *B*=-0.01 $CI_{90\%}$ = [-0.04, 0.02] | ***B*=0.07 $CI_{99.9\%}$ = [0.02, 0.12]** | *B*=-0.00 $CI_{90\%}$ = [-0.03, 0.02] |
| vStriatum | *B*=-0.03 $CI_{90\%}$ = [-0.06, 0.01] | *B*=-0.01 $CI_{90\%}$ = [-0.04, 0.03] | *B*=0.02 $CI_{90\%}$ = [-0.01, 0.06] | *B*=0.01 $CI_{90\%}$ = [-0.02, 0.04] | ***B*=0.03 $CI_{90\%}$ = [0.00, 0.07]** | *B*=0.01 $CI_{90\%}$ = [-0.03, 0.04] | *B*=-0.00 $CI_{90\%}$ = [-0.03, 0.03] |
| vmPFC | *B*=-0.03 $CI_{90\%}$ = [-0.06, 0.00] | *B*=-0.03 $CI_{90\%}$ = [-0.06, 0.00] | *B*=-0.00 $CI_{90\%}$ = [-0.04, 0.03] | *B*=0.00 $CI_{90\%}$ = [-0.03, 0.03] | ***B*=0.04 $CI_{90\%}$ = [0.00, 0.07]** | *B*=-0.03 $CI_{90\%}$ = [-0.07, 0.00] | *B*=0.00 $CI_{90\%}$ = [-0.03, 0.03] |

aInsula = anterior insula, BNST = bed nucleus of the stria terminalis, dACC = dorsal anterior cingulate cortex, PAG = periaqueductal gray, vStriatum = ventral striatum, vmPFC = ventromedial prefrontal cortex. Effects that reached (marginal) significance (i.e., ≥90% posterior credible interval was non-overlapping with zero) are displayed in bold.

**Table S21. Post-hoc comparisons for the interaction between reward and decision: reward effect in the *outcome anticipation phase*, separated by decision (approach vs avoid).**

| | *Reward effect* | |
|---|---|---|
| | Approach | Avoid |
| vStriatum | $B$=0.01<br>$CI_{90\%}$ = [-0.03, 0.05] | ***B*=-0.06**<br>**$CI_{90\%}$ = [-0.11, -0.00]** |
| vmPFC | $B$=0.01<br>$CI_{90\%}$ = [-0.03, 0.05] | ***B*=-0.06**<br>**$CI_{90\%}$ = [-0.11, -0.01]** |

vStriatum = ventral striatum, vmPFC = ventromedial prefrontal cortex. Effects that reached (marginal) significance (i.e., ≥90% posterior credible interval was non-overlapping with zero) are displayed in bold.

**Table S22. Post-hoc comparisons for the interaction between threat and decision: threat effect in the *outcome anticipation phase*, separated by decision (approach vs avoid).**

| | *Threat effect* | |
|---|---|---|
| | Approach | Avoid |
| aInsula | ***B*=0.16**<br>**$CI_{99.9\%}$ = [0.07, 0.26]** | $B$=-0.01<br>$CI_{90\%}$ = [-0.08, 0.05] |
| BNST | ***B*=0.08**<br>**$CI_{99\%}$ = [0.01, 0.15]** | $B$=-0.04<br>$CI_{90\%}$ = [-0.11, 0.02] |
| dACC | ***B*=0.09**<br>**$CI_{99.9\%}$ = [0.03, 0.15]** | $B$=-0.04<br>$CI_{90\%}$ = [-0.08, 0.01] |
| PAG | ***B*=0.14**<br>**$CI_{99.9\%}$ = [0.04, 0.24]** | $B$=0.02<br>$CI_{90\%}$ = [-0.06, 0.09] |
| Thalamus | ***B*=0.08**<br>**$CI_{99.9\%}$ = [0.02, 0.14]** | ***B*=0.05**<br>**$CI_{95\%}$ =[-0.10, -0.00]** |

aInsula = anterior insula, BNST = bed nucleus of the stria terminalis, dACC = dorsal anterior cingulate cortex, PAG = periaqueductal gray. Effects that reached (marginal) significance (i.e., ≥90% posterior credible interval was non-overlapping with zero) are displayed in bold.

**Table S23. Post-hoc comparisons for the interaction between reward, threat, and decision: differences in threat effect between approach and avoid decisions in the *outcome anticipation phase*, separated for low and high reward conditions.**

| | *Differences in threat effect between approach and avoidance decisions* | |
|---|---|---|
| | Low reward | High reward |
| aInsula | ***B*=0.28**<br>**$CI_{99.9\%}$ = [0.07, 0.50]** | $B$=0.07<br>$CI_{90\%}$ = [-0.06, 0.19] |

aInsula = anterior insula. Effects that reached (marginal) significance (i.e., ≥90% posterior credible interval was non-overlapping with zero) are displayed in bold.

**Table S24. Post-hoc comparisons: threat effect in low reward conditions in the *outcome anticipation phase*, separated by decision (approach vs avoid).**

| | *Threat effect in low reward conditions* | |
|---|---|---|
| | Approach | Avoid |
| aInsula | ***B*=0.20**<br>**$CI_{99.9\%}$ = [0.04, 0.36]** | ***B*=-0.09**<br>**$CI_{90\%}$ = [-0.16, -0.01]** |

aInsula = anterior insula. Effects that reached (marginal) significance (i.e., ≥90% posterior credible interval was non-overlapping with zero) are displayed in bold.
